# Brief illumination of an optoGPCR elicits prolonged inhibition of *Drosophila* behavior

**DOI:** 10.64898/2026.05.26.727874

**Authors:** Nicole Mynyi Lee, Yishan Mai, Leif Dalberg, Jonathan Charles Anns, Danesha Devini Suresh, Zhiyi Zhang, Adam Claridge-Chang

**Author notes:** Equal contribution (Nicole Mynyi Lee, Yishan Mai).

## Abstract

Light-actuated G protein–coupled receptors (optoGPCRs) are a class of opsins that act through intracellular signaling cascades. As optogenetic tools, bistable inhibitory optoGPCRs are attractive as they decouple the duration of silencing from the duration of illumination: brief light pulses drive prolonged signaling. Opsin3 from the mosquito *Anopheles stephensi* (MosOPN3) is a bistable Gα_i/o_-coupled opsin that produces sustained inhibition in mammalian neurons and *C. elegans*, but its utility in vinegar flies, *Drosophila melanogaster*, remains untested. Here we tested three optoGPCRs in adult flies; MosOPN3 emerged as the most effective and was benchmarked against an established inhibitor, *Guillardia theta* anion channelrhodopsin 1 (GtACR1), in walking and memory assays.

MosOPN3 inhibitory potency was comparable to that of GtACR1 across most neuronal classes tested, with substantially lower developmental toxicity. Rearing flies on all-*trans*-retinal–supplemented food throughout development improved both viability and inhibitory efficacy for both opsins. A single pulse of green light (as brief as 5 s) inhibited behavior for minutes, and was sufficient to ablate aversive olfactory learning over a one-minute training epoch. For flies, MosOPN3 is a low-toxicity optogenetic inhibitor for flies that yields minutes of silencing from brief light pulses—especially useful where continuous actuating light would be confounding.

**Key points:**

- To our knowledge, *Anopheles stephensi* Opsin3 (OPN3) is the first GPCR opsin that inhibits behavior in adult *Drosophila melanogaster*.
- OPN3 has inhibitory efficacy close to that of GtACR1, with superior viability.
- Brief light pulses drive sustained inhibition lasting minutes, eliminating the need for continuous illumination.
- For both GtACR1 and OPN3, rearing flies on ATR-supplemented food enhances inhibition efficacy and improves developmental viability.

## Introduction

Optogenetic tools, which enable the rapid and precise control of cellular physiology using light, have emerged as a critical tool for studying circuit function (Boyden et al., 2005; Deisseroth, 2015; Emiliani et al., 2022; Rost et al., 2022; Yizhar et al., 2011; Zemelman et al., 2002). The most widely used optogenetic tools are the cation channelrhodopsins (ChR), algae-derived light-gated channels, which typically cause membrane depolarization and thus neuronal activation (Boyden et al., 2005; Deisseroth, 2015; Li et al., 2005; Nagel et al., 2003, 2002; Yizhar et al., 2011; Zemelman et al., 2002). In contrast, the development of high-efficacy inhibitory optogenetic tools has been more challenging (Berndt et al., 2014; Chow et al., 2010; Govorunova et al., 2015; Gradinaru et al., 2007; Han and Boyden, 2007; Wietek et al., 2014; Yizhar et al., 2011), with earlier tools requiring high expression and strong light intensities (Gradinaru et al., 2010, 2008; Mahn et al., 2016), limiting their utility in vivo.

Anion channelrhodopsins (ACRs) (Berndt et al., 2014; Govorunova et al., 2015; Mohammad et al., 2017; Wietek et al., 2014) and kalium channelrhodopsins (KCRs) (Govorunova et al., 2022; Ott et al., 2024; Vierock et al., 2022) are powerful tools for optogenetic silencing.

However, even these methods have some limitations. ACR action depends on the chloride gradient: in compartments with elevated intracellular chloride such as axonal terminals, non-neuronal cells, and immature neurons (Duebel et al., 2006; Kaila et al., 2014; Kilb, 2021; Mueller et al., 1983; Price and Trussell, 2006), actuation can drive depolarization and excitation rather than inhibition (Himmel et al., 2023; Mahn et al., 2018, 2016; Malyshev et al., 2017; Messier et al., 2018; Szabadics et al., 2006). Prolonged chloride conductances can also lead to secondary effects such as potassium redistribution, introducing further unpredictability in neuronal excitability (Kaila et al., 2014; Parrish et al., 2023). Similarly, KCR variants can have excitatory activity, particularly during longer illumination epochs (Duan et al., 2026; Govorunova et al., 2022; Vierock et al., 2022). Extant ACR and KCR channels close in the dark, meaning that inhibition lasts only as long as illumination, and that prolonged silencing thus requires prolonged illumination. This constrains experimental designs requiring sustained or repeated pulsed illumination, thereby increasing cumulative light exposure over long-duration experiments, and complicating compatibility with concurrent calcium imaging or experiments where continuous light affects behavior and/or system physiology. Such limitations have long motivated interest in bistable optogenetic tools whose action outlasts the light pulse (Berndt et al., 2009; Koyanagi et al., 2004); these could allow prolonged inhibition with only brief light actuation (Mahn et al., 2021; Rodriguez-Rozada et al., 2022; Wada et al., 2026; Wietek et al., 2024, 2017).

Light-actuated G-protein-coupled receptors (‘optoGPCRs’) are part of the optogenetic toolkit for manipulating neuronal physiology (Marcus and Bruchas, 2023; Spangler and Bruchas, 2017). Distinct from channelrhodopsins, which directly conduct ion flow, optoGPCRs act through intracellular signaling cascades, similar to other GPCR-mediated processes triggered by neuropeptides and neuromodulators (Huang and Thathiah, 2015). A variety of GPCR opsins have been developed for optogenetic interventions. The original optogenetic tool was chARGe, an optoGPCR method that co-expressed an insect rhodopsin with two signaling components (Zemelman et al., 2002). Engineering different classes of GPCRs enabled the development of opto-XRs, hybrid proteins with light-activated control of specific guanine nucleotide-binding proteins (G proteins e.g. Gα_s_, Gα_i/o_, or Gα_q_) (Airan et al., 2009; Kim et al., 2005). JellyOp (from the jellyfish *Carybdea rastonii*) enables light actuation of the Gα_s_ pathway (Bailes et al., 2012; Koyanagi et al., 2008), while lamprey parapinopsin (LamPP, from *Lethenteron camtschaticum*) and mosquito Opsin3 (MosOPN3 *Anopheles stephensi*) couple to Gα_i/o_ and suppress neuronal activity through inhibition of adenylyl cyclase and reducing neuronal excitability in mammals (Copits et al., 2021; Hagio et al., 2023; Koyanagi et al., 2013, 2004; Mahn et al., 2021; Rodgers et al., 2021) and *Caenorhabditis elegans* (Koyanagi et al., 2022, 2013). These optoGPCRs allow light control of neuromodulatory cascades and neuronal activity using the cell’s native G-protein signaling machinery, which represents a distinct advantage over the exogenous expression of potentially toxic channelrhodopsins (Luo et al., 2020; Marcus and Bruchas, 2023; Sakami et al., 2011).

Among these, MosOPN3 is a promising candidate due to its bistability and robust inhibitory effects (Koyanagi et al., 2022, 2013; Mahn et al., 2021). MosOPN3 has shown coupling to inhibitory Gα_i/o_ proteins in multiple binding and activity assays (Koyanagi et al., 2022, 2013; Mahn et al., 2021; Wietek et al., 2024). In mice, a trafficking-enhanced variant, enhanced OPN3 (eOPN3), was found to suppress synaptic transmission by modulating voltage-gated calcium channels at presynaptic sites (**Figure 1B**, (Koyanagi et al., 2022, 2013; Mahn et al., 2021), successfully inhibiting hippocampal, cortical, thalamic, and mesencephalic circuits (Mahn et al., 2021).

**Figure 1.**
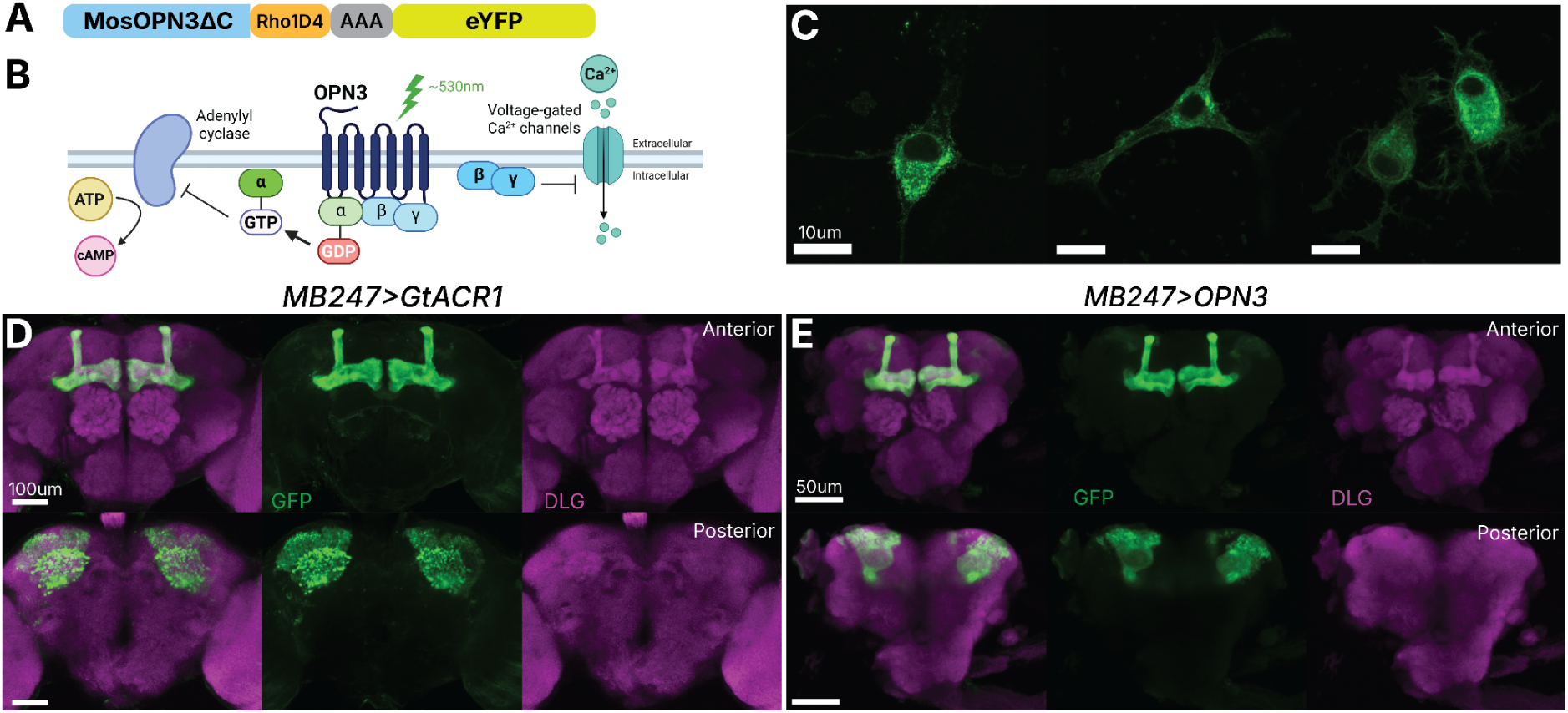
OPN3 expression and localization in mouse N2a cells and *Drosophila* mushroom-body neurons (A) Schematic of fly codon-optimized MosOpn3ΔC construct with Rho1D4 epitope tag (orange), AAA linker (gray) and eYFP reporter used in this study. **(B)** Mechanism of MosOPN3 action: upon green-light activation, OPN3 is thought to couple to endogenous Gα_i/o_ proteins. Gβγ subunits modulate presynaptic voltage-gated Ca²⁺ channels, while Gα_i_ inhibits adenylyl cyclase, reducing cAMP levels. **(C)** Representative images of OPN3-eYFP expression in transfected mouse N2a cells stained with anti-GFP antibody. Scale bars = 10 µm. n = 4 **(D, E)** Representative confocal images of adult *Drosophila* brains expressing GtACR1 (D) or OPN3 (E) under *MB247-GAL4* driver. Images show anterior and posterior views with GFP (green) and Anti-Discs large (DLG) neuropil marker (magenta). Scale bars for GtACR1 = 100 µm; OPN3 = 50 µm. See **Supplementary Figure 4** for summed pixel intensity quantification.

To date, for the important neurogenetic system the adult vinegar fly (*Drosophila melanogaster*), no inhibitory optoGPCR has been demonstrated to produce behavioral inhibition. In this study, we evaluate the inhibitory potency of several bistable optoGPCRs in flies. One variant, a C-terminally truncated MosOPN3 (MosOpn3ΔC) without additional trafficking signals was found to be an effective inhibitor. We quantified its ability to suppress behavior, benchmarked its efficiency against the established optogenetic inhibitor GtACR1, and assessed its suitability for short-pulse inhibition. Our results establish MosOpn3ΔC as an advantageous inhibitory optogenetic tool that demonstrates potent inhibition, reduced toxicity compared to GtACR1, and prolonged effects with only brief pulses of actuating illumination.

## Results

### In adult *Drosophila*, OPN3 is an effective inhibitory optoGPCR

To identify an effective inhibitory optoGPCR for adult *Drosophila*, we screened three Gα_i/o_-coupled candidates in a climbing assay: *Platynereis dumerilii* ciliary opsin (PdCO), MosOPN3ΔC, and its trafficking-enhanced variant, eOPN3 (overview of opsin constructs in **Supplementary Figure 1**). PdCO has been shown to inhibit neurons in mice, *Caenorhabditis elegans*, and *Drosophila* larvae (Wietek et al., 2024), but has not been reported for adult flies. eOPN3, which incorporates a Kir2.1-derived membrane trafficking signal and ER export motif into MosOPN3ΔC, has been shown to suppress synaptic transmission in mammalian neurons and induce an ipsiversive rotational bias in mice (Mahn et al., 2021).

Each MosOPN3-based candidate opsin gene was synthesized with codon usage optimized for *Drosophila* with fusion tags and linkers (see Methods).

Of these three candidates tested in adult *Drosophila*, only the fly-codon optimized version of MosOpn3ΔC with protein tags (henceforth referred to as ‘OPN3’; **Figure 1A**) produced robust behavioral inhibition. PdCO produced little to no behavioral inhibition in the adult flies (**Supplementary Figure 2**). The fly codon-optimized eOPN3 also did not consistently produce robust inhibition (**Supplementary Figure 3**), despite the strong precedents (Mahn et al., 2021) and prior work showing that trafficking and export motifs (Ma et al., 2001; Stockklausner et al., 2001; Stockklausner and Klocker, 2003) support efficient opsin expression in multiple systems *(Gradinaru et al., 2010; Mahn et al., 2021; Ott et al., 2024)*. We therefore focused on OPN3 (**Figure 1A**) for further characterization.

### In *Drosophila* neurons, OPN3 localizes to distal neurites efficiently

To characterize OPN3 (**Figure 1A**) expression and localization, we transfected mouse Neuro-2a (N2a) neuroblastoma cells with the construct and observed robust yellow fluorescent protein (YFP) signals 48 h post-transfection. Transfected cells displayed a predominantly intracellular distribution pattern (**Figure 1C**), contrasting with the improved plasma-membrane localization achieved with eOPN3 (Mahn et al., 2021). MosOpn3 nevertheless was seen to confer light sensitivity to cultured mammalian cells (Koyanagi et al., 2013), suggesting that membrane trafficking signals are not essential for basic functionality.

To benchmark OPN3 against GtACR1, a widely adopted inhibitory optogenetic tool in *Drosophila* (Mohammad et al., 2017), we expressed both constructs using the MB247-GAL4 driver line targeting mushroom body neurons. Confocal imaging revealed strong YFP fluorescence in both anterior (α/β lobes) and posterior (cell bodies and calyces) brain regions (**Figure 1D–E**). GtACR1 showed a posterior bias in summed pixel intensity across slices (Hedges’ *g* = –0.81), whereas OPN3 exhibited a more balanced distribution across brain compartments, with slightly elevated anterior signal (Hedges’ *g* = +0.23), indicating more efficient trafficking into distal neurites (**Supplementary Figure 4)**.

### OPN3 shows robust developmental viability

A practical concern when characterizing optogenetic tools is developmental toxicity. To assess whether OPN3 expression impairs *Drosophila* development, OPN3 lines were crossed with a panel of GAL4 drivers representing different neuronal populations: pan-neuronal lines (*elav-Gal4*, *nSyb-Gal4*), glutamatergic (*vGlut-Gal4,* ∼17% of brain neurons, and *OK371-Gal4*), GABA-ergic (*vGAT-Gal4,* ∼10% of brain neurons) (Amin et al., 2023; Croset et al., 2018; Daniels et al., 2008; Eckstein et al., 2024) and *GMR59D01-GAL4* (henceforth referred to as *jus-Gal4*), a driver derived from the bang-sensitive seizure gene *julius seizure* that labels central neurons including the optic lobe and descending populations (Horne et al., 2017; Jenett et al., 2012; Pfeiffer et al., 2010). Crosses were maintained under standard rearing conditions and offspring were scored for eclosion.

We found that crosses of OPN3 with the selected driver lines were all viable, while five of six drivers with GtACR1 were inviable on standard food (**Figure 2C**). Quantification of lethality proportions revealed that *elav>OPN3* flies showed eclosion failure rates (Δlethality = +0.46) comparable to control *elav-Gal4/+* levels (Δlethality = +0.32; **Figure 2D**), indicating that pan-neuronal OPN3 expression imposes only a modest additional developmental cost beyond background. On the other hand, *elav>GtACR1* progeny were entirely inviable when reared on standard food (Δlethality proportion = +1.00; **Figure 2D**) with pupae displaying characteristic melanization indicative of developmental failure. These results demonstrate that, even when expressed with broad neuronal drivers, OPN3 has only a minor effect on developmental viability.

**Figure 2.**
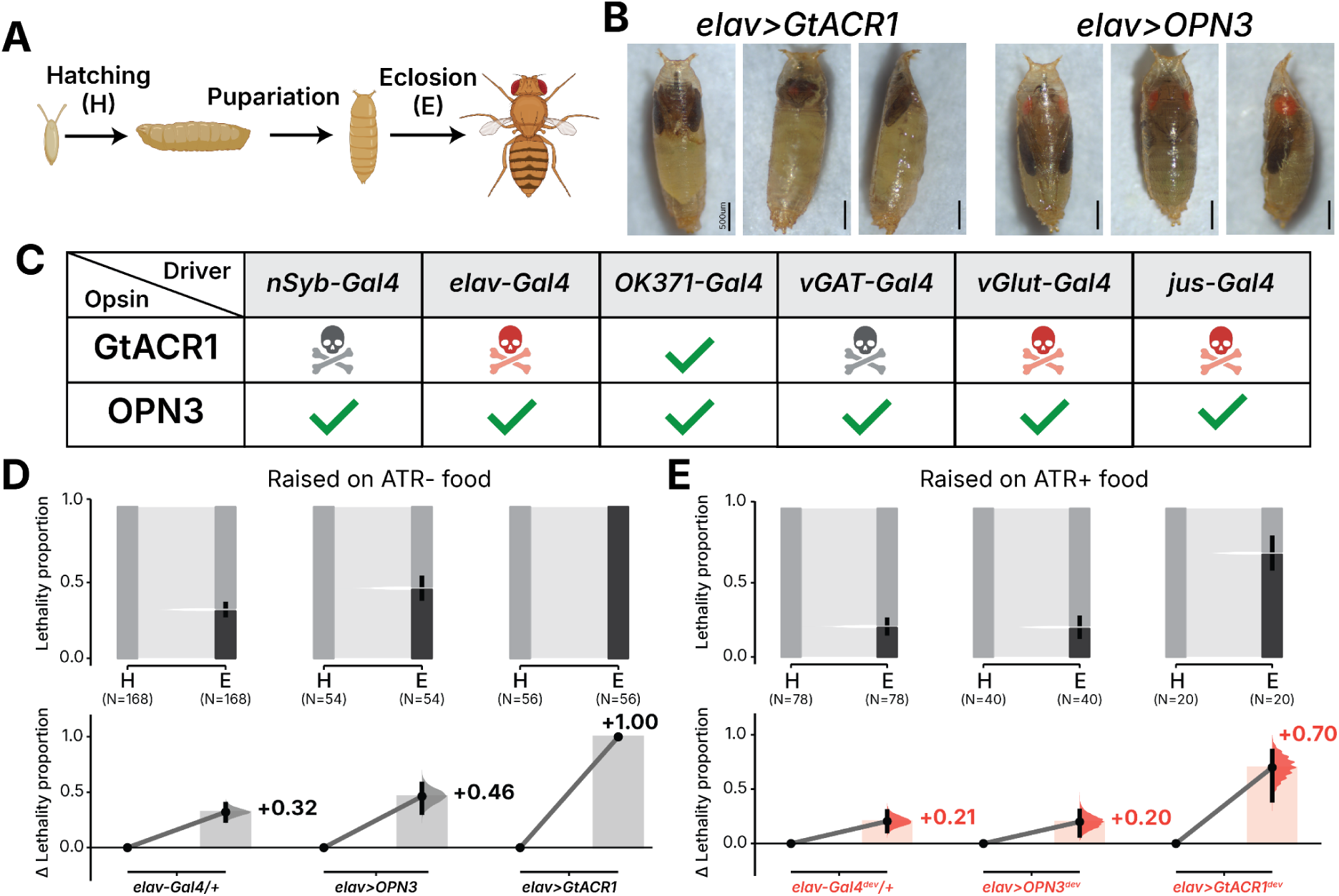
OPN3 does not impair *Drosophila* developmental viability and ATR supplementation rescues GtACR1 developmental toxicity. (A) Schematic of the *Drosophila melanogaster* life cycle showing developmental stages. Hatching (H) from the embryo proceeds through larval stages to pupariation, followed by pupal development and eclosion (E) to the adult stage. **(B)** Representative images of pupae expressing GtACR1 (left) or OPN3 (right) under the *elav-GAL4* driver, demonstrating melanization. Scale bars = 500um. **(C)** Summary table showing driver-opsin combinations tested for viability. Black skulls indicate lethality regardless of diet, red skulls indicate lethality only when raised on non ATR food; checkmark indicates successful development to adulthood on both non ATR and ATR food. **(D, E)** Proportion plots quantifying developmental lethality for each driver-opsin combination reared on **(D)** regular food or **(E)** ATR supplemented food, assessed at developmental stage H and E. Gray bars represent the proportion of individuals that successfully completed each developmental stage; black bars represent the proportion that failed. Sample sizes (N) indicated below each bar. Error bars represent 95% CI. Superscripted ‘dev’ indicates flies were raised on ATR throughout development and post eclosion.

### Developmental ATR supplementation of opsin flies improves viability

Our experiments used ATR feed supplementation, based on the idea that 13-*cis* retinal forms from thermally isomerized dietary ATR and would be sufficient to bind the apo-opsins in vivo and form functional photopigments (Koyanagi et al., 2013). In earlier experiments, we raised flies on standard food during development and transferred them to

ATR-supplemented food for 2–3 d post-eclosion (sATR). We therefore tested whether providing ATR throughout development would mitigate this lethality. Indeed, providing flies with ATR-supplemented food throughout development and adulthood (∼14–17 d total; devATR) partially relieved developmental lethality. In *elav>GtACR1, devA*TR reduced the lethality from 100% to 70%, although still substantially higher than both controls (**Figure 2D, E**). *elav>OPN3* similarly shows improvement in lethality score from 46% to 20% (**Figure 2D, E**). Complete counts for hatching and eclosion across all genotypes are provided in Supplementary Data.

### OPN3 produces robust and reversible behavioral inhibition

Under sATR conditions, we tested OPN3 flies on a climbing assay that quantifies negative geotaxis (**Figure 3A**) (Ganetzky and Flanagan, 1978; Jones and Grotewiel, 2011; Le Bourg and Lints, 1992; Miquel et al., 1972; Riemensperger et al., 2013; Sun et al., 2018) to assess its efficacy as an inhibitory optogenetic tool. Upon green illumination (530 nm, 22 μW/mm^2^), *elav>OPN3* flies showed suppression of climbing height, while genetic controls showed no light-dependent changes (Δg = −2.56, **Figure 3B, E, Supplementary Figure 5A**). OPN3 produced robust inhibition across nearly all the broad neuronal populations targeted by the drivers listed in Figure 2C, with large effects on both climbing height (**Figure 3E**) and speed (**Supplementary Figure 6**). *OK371-Gal4* was the sole driver tested that failed to produce a behavioral effect with OPN3, with flies showing no inhibition (Height Δg = −0.04, **Supplementary Figure 7E, G**). Complete effect sizes with 95% CIs for all genotypes across all conditions (genetic controls, light status, food status) are provided in <u>Supplementary Data</u>.

**Figure 3.**
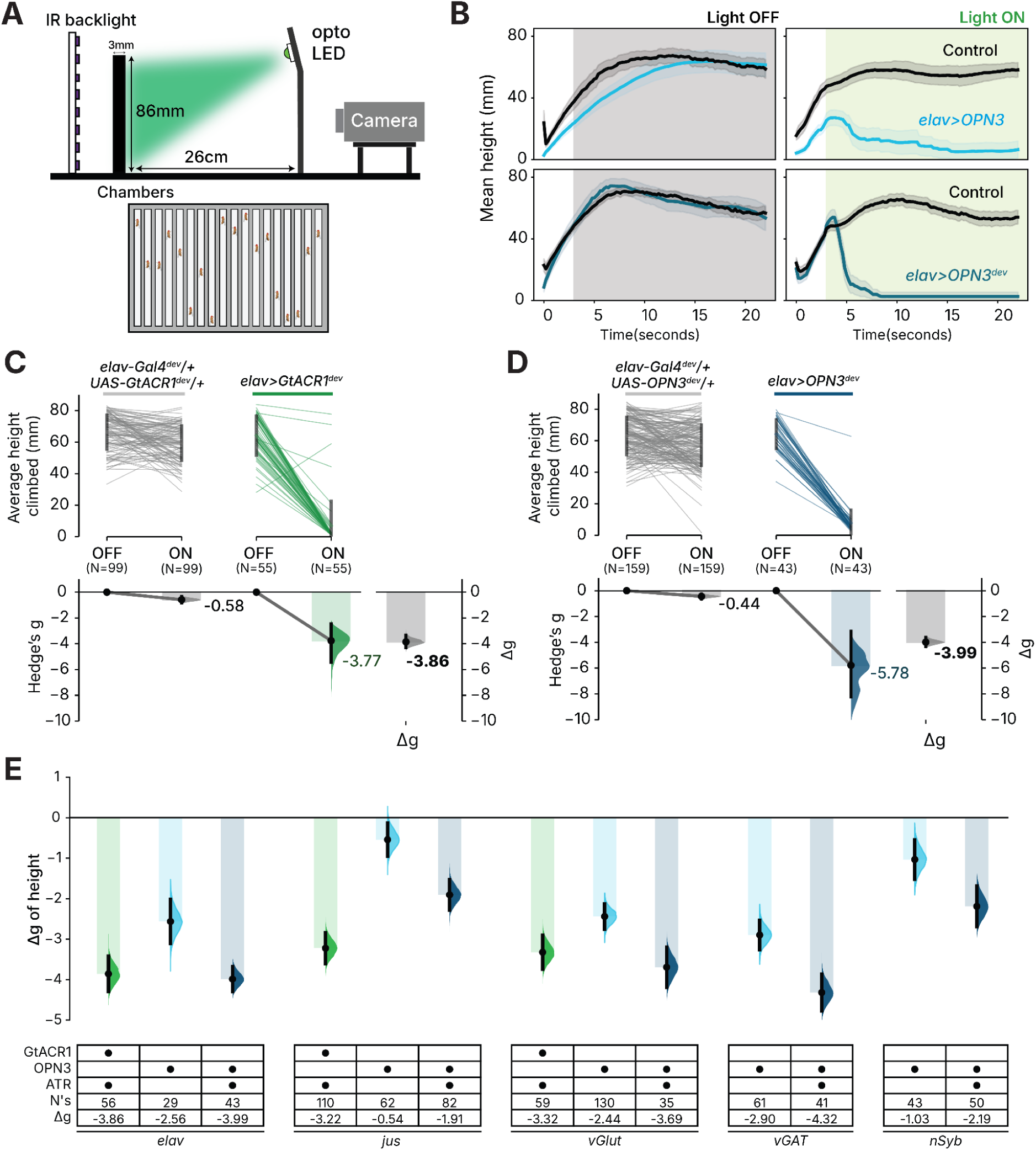
Across neuronal classes, OPN3 inhibition is comparable to GtACR1 (A) Schematic of climbing assay apparatus. Flies are loaded into vertical chambers (86 mm), agitated to the bottom, and recorded via infrared backlight and camera during baseline and optogenetic stimulation (530 nm). **(B)** Representative traces of climbing height for *elav>OPN3* flies on sATR (top) and devATR (bottom) compared to the driver (*elav-Gal4/+*) and responder (*UAS-OPN3*) controls (black) matched to the same dietary ATR protocol. Gray background, light OFF; green background, light ON. **(C, D)** Estimation plot of climbing height for (**C**) *elav>GtACR1*^dev^ and (**D**) *elav>OPN3*^dev^ across light OFF and light ON conditions fed on ATR supplemented food. Top: Raw data; error bars indicate SD. Bottom: Effect sizes; error bars indicate 95% CI. Right bottom: Δg effect size. **(E)** Summary of Δg height effect sizes across multiple neuronal drivers (*elav-Gal4*, *jus-Gal4*, *vGAT-Gal4*, *vGlut-Gal4* and *nSyb-Gal4*) for GtACR1 (green) and OPN3 (blue). Gridkey indicates opsin, ATR supplementation, sample size (N’s) and Δg effect size. Error bars represent 95% CI. Darker colored markers indicate devATR; lighter markers indicate sATR. Green, GtACR1; blue, OPN3. Superscripted ‘dev’ indicates flies were raised on ATR throughout development and post eclosion.

### Developmental ATR supplementation enhances opsins’ performance

The comparisons above were conducted under different dietary conditions: GtACR1^dev^ flies received devATR, while OPN3 flies received sATR. To enable direct comparison, we raised OPN3 crosses under devATR (OPN3^dev^), matching GtACR1^dev^ dietary conditions. ATR supplementation approximately doubled the inhibitory Δg effect size on climbing height for OPN3^dev^ compared to OPN3 on standard food across most drivers tested (**Figure 3E**). A similar pattern was observed for speed, though with smaller absolute differences between conditions (**Supplementary Figure 6**). *OK371>OPN3*^dev^ remained ineffective (Δg = −0.23; **Supplementary Figure 7**), consistent with results on standard food. Under matched dietary conditions, OPN3^dev^ produced effect sizes comparable to GtACR1^dev^ across most responsive driver lines (**Figure 3E**), indicating that ATR-supplemented OPN3 achieves inhibitory efficacy equivalent to the current benchmarked tool.

### Actuation of OPN3 in memory cells during training robustly inhibits learning

As the previous drivers used represented broad neuronal populations, we decided to examine the efficacy of OPN3 to inhibit behaviors that rely on small groups of neurons. For this, we used three drivers: *Th-Gal4*, MB320C split-Gal4 combination, and *MB247-Gal4 (Aso et al., 2014; Dubnau et al., 2001; Mao and Davis, 2009; Zars et al., 2000)*. *Th-Gal4* drives expression in the dopaminergic system, capturing ∼130 cells per hemisphere (Claridge-Chang et al., 2009; Mao and Davis, 2009). The specific MB320C split-Gal4 driver drives in a single dopaminergic neuron per hemisphere: the PPL101 cell, also known as PPL1-γ1pedc (Aso et al., 2012; Pfeiffer et al., 2010). As inhibition of either the *Th-Gal4* or PPL101 dopamine neurons during training causes defective aversive associative memory (Aso et al., 2012; Aso and Rubin, 2016; Berry et al., 2012; Otto et al., 2020), we used an olfactory learning paradigm to characterize the efficacy of OPN3 inhibition (**Figure 4A–B**; see Methods). Similarly, this paradigm allowed us to examine OPN3 efficacy in the *MB247-Gal4* subset of mushroom body-intrinsic Kenyon cells (KCs), the inhibition of which also blocks learning (de Belle and Heisenberg, 1994; Heisenberg et al., 1985; Turner et al., 2008; Zars et al., 2000).

**Figure 4.**
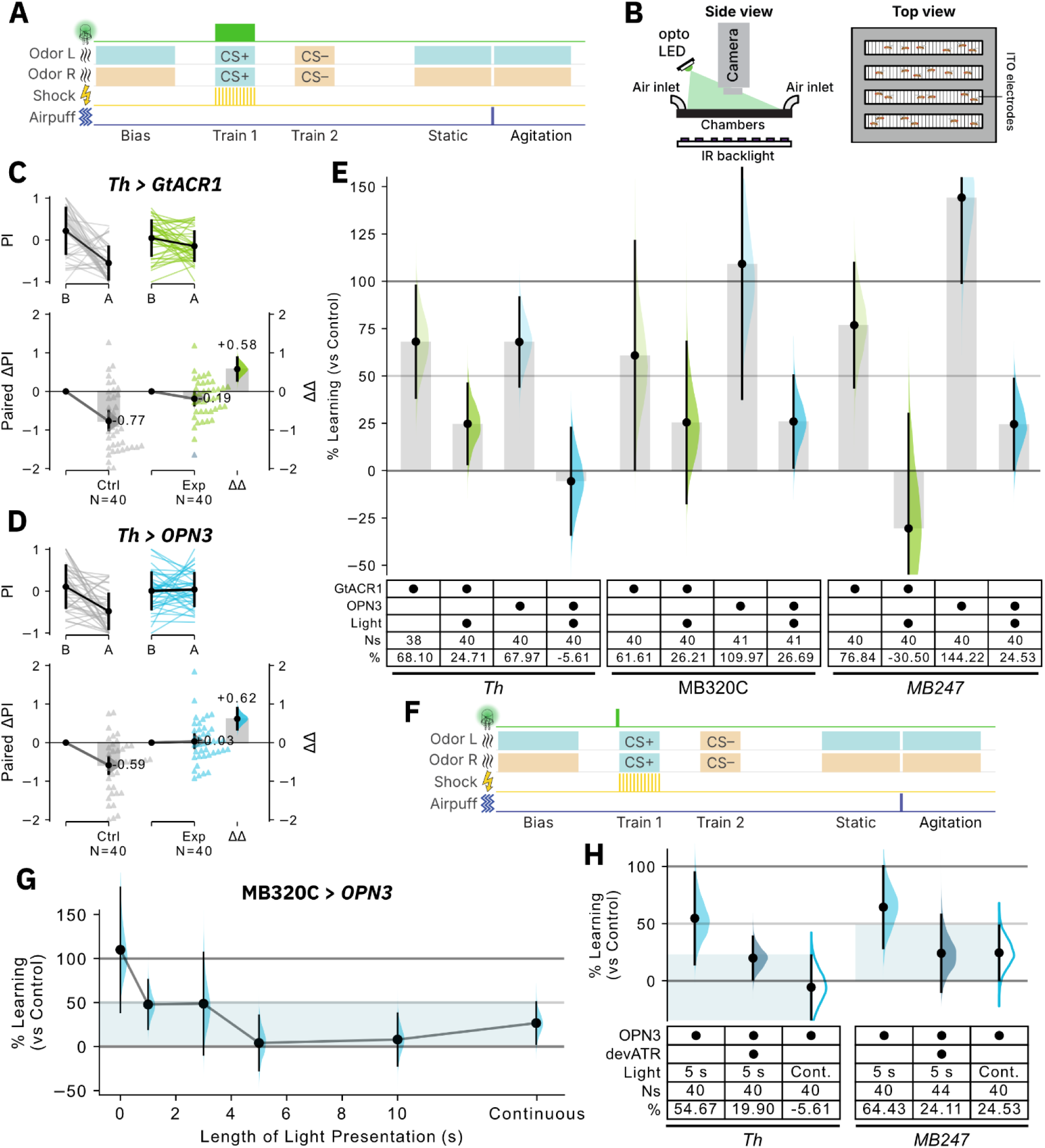
OPN3 inhibits dopaminergic neurons and Kenyon cells during shock training. (A) Schematic of the experimental protocol used for panels 4C–E and Supplementary Figure 8. For light-on experiments, green light is presented throughout the 1-min shock training epoch; green light is omitted for light-off. **(B)** Schematic of the Multifly Olfactory Trainer. **(C–D)** Broad inhibition of dopaminergic neurons with either GtACR1 or OPN3 dramatically reduced aversive learning. Top: Raw data; error bars indicate SD. Bottom: Effect sizes; error bars indicate 95% CI. **(E)** Summary plot of % learning for *Th>GtACR1, Th>OPN3, MB320C>GtACR1, MB320C>OPN3*, *MB247>GtACR1, MB247>OPN3* comparing light-off and light-on conditions. “Controls” refers to genetic controls (i.e. pooled driver-only and responder-only controls) for each respective genotype. Raw data from Supplementary Figure 8A-J. Error bars indicate 95% CI. Some CIs have been truncated for readability; refer to Supplementary Data for full CIs. **(F)** Schematic of the experimental protocol used for panel G and Supplementary Figures 9-10. A short pulse of light (ranging from 1 s to 10 s) was presented directly before the shock epoch. **(G)** 5 s and 10 s light pulses resulted in inhibition greater than continuous light presentation in *MB320C>OPN3* flies. 0 s and continuous light are identical to light-off and light-on data from Supplementary Figure 8E–F. Shaded region: 95% CI of continuous light-on condition. **(H)** ATR supplementation improved inhibition with 5 s light pulse in *Th>OPN3* and *MB247>OPN3* flies to a level similar to continuous light inhibition (“Cont.”) without ATR supplementation. “Controls” refers to genetic controls. Raw data from Supplementary Figure 9G,O for 5 s light, and Figure 4D for continuous light for *Th>OPN3*; Supplementary Figure 9K,P for 5 s light and Supplementary Figure 8J for continuous light for *MB247>OPN3*. Note that % learning for continuous light is also plotted in panel E. Shaded regions: 95% CI of continuous light-on condition for each genotype. Means and CIs for all panels are available in Supplementary Data.

For all three drivers, we compared OPN3 to GtACR1 by actuating with green light for the full duration of the odor–shock training epoch (**Figure 4A**). We assessed the change in preference index (ΔPI) between the pre-training bias test period and post-training agitation test periods and compared the ΔPI between control and test flies to obtain a ΔΔPI (see Methods). As expected, illumination during training resulted in poor learning for the experimental flies (ΔPI = –0.19 and +0.03 respectively). Compared to uninhibited controls, GtACR1 and OPN3 actuation each resulted in large avoidance-memory deficits, ΔΔPI = +0.58 and +0.62 respectively, with the large, positively signed effect sizes indicating strong impairments of aversive memory (**Figure 4C–D**). We further quantified impairment of learning by calculating a % learning metric compared to genetic controls (see Methods), and confirmed that both *Th>GtACR1* and *Th>OPN3* flies showed greatly impaired memory compared to controls; the fractions of control learning were just 24.71% and –5.61% respectively (**Figure 4E**). In comparison, when light was not presented during training, both *Th-Gal4>GtACR1* and *Th-Gal4>OPN3* learned normally, although with weaker ΔPIs compared to controls, reflected as a slightly positive ΔΔPI (ΔPI =-0.48 and-0.47; ΔΔPIs = +0.22 and +0.22; % learning: 68.02% and 67.73%; **Figure 4E, Supplementary Figure 8A–B**). These dark effects in opsin-expressing flies were likely due to non-actuated activity and/or toxicity from opsin expression. *MB320C>GtACR1, MB320C*>*OPN3*, *MB247>GtACR1* and *MB247>OPN3* flies all also showed learning impairments when light was presented during the shock epoch. Expression of GtACR1 alone resulted in reductions in learning even without illumination in all three drivers; however in MB320C and MB247, expression of OPN3 in the dark did not result in learning reductions (**Figure 4E**), thus indicating that OPN3 has a lower effect on baseline cell function when not actuated compared to GtACR1.

### Brief actuation of OPN3 in specific circuits inhibits memory formation

We next tested whether actuation of OPN3 with short pulses of light would also lead to sustained inhibition. Using the same associative memory assay, we used different durations of light directly prior to shock to test whether it was possible to inhibit dopaminergic neurons or KCs for the full 1-min training period (**Figure 4F**). Prolonged inhibition with OPN3 was most pronounced in MB320C>*OPN3* flies, where 5 s and 10 s of light illumination were sufficient to ablate learning (% learning: 4.10% and 7.98% respectively; **Figure 4G**, **Supplementary Figure 9A–D**). In *Th>OPN3* flies, learning was greatly reduced with 10 s of light stimulation (**Supplementary Figure 9E–H,M**), while in *MB247>OPN3* flies, learning was greatly reduced at 3 s and 10 s (**Supplementary Figure 9I–L,N**). In contrast, when GtACR1 was used, short pulses of light failed to induce reduction in learning with all drivers and with all times tested (**Supplementary Figure 10**).

As 5 s of light presentation failed to reduce learning below 50% in *Th>OPN3* and *MB247>OPN3* flies, we decided to test whether developmental ATR supplementation would improve the long-term inhibitory effect induced in these conditions, as previously demonstrated with broad drivers (**Figure 3E**). We found that developmental ATR supplementation indeed further reduced learning in both drivers, resulting in effect sizes similar to that of continuous light presentation in flies without developmental ATR supplementation (*Th>OPN3^dev^,* % learning with 5 s light = 19.90% compared to *Th>OPN3*, % learning in continuous light = 19.99%; *MB247>OPN3^dev^,* % learning with 5 s light = 24.11%, *MB247>OPN3*, % learning with continuous light = 24.53%; **Figure 4H, Supplementary Figure 9O–P**).

### Brief actuation of OPN3 elicits a prolonged inhibition of behavior

Based on the improved efficacy observed with developmental ATR supplementation, all subsequent experiments reared OPN3 crosses on developmentally ATR-supplemented food (OPN3^dev^). In climbing assays, it was observed that OPN3-expressing flies failed to resume walking after light offset, whereas GtACR1-expressing flies recovered immediately. This persistence is consistent with the bistability of OPN3 where it retains its retinal chromophore after photoactivation, enabling sustained signaling following brief light pulses (Koyanagi et al., 2013; Mahn et al., 2021). To characterize this prolonged inhibition, we used the Trumelan assay for continuous behavioral monitoring over extended periods (**Figure 5A**). Speed recordings confirmed that GtACR1-mediated inhibition terminated with light offset, while OPN3-mediated inhibition persisted for several minutes afterward, regardless of ATR supplementation status (**Figure 5B**).

**Figure 5.**
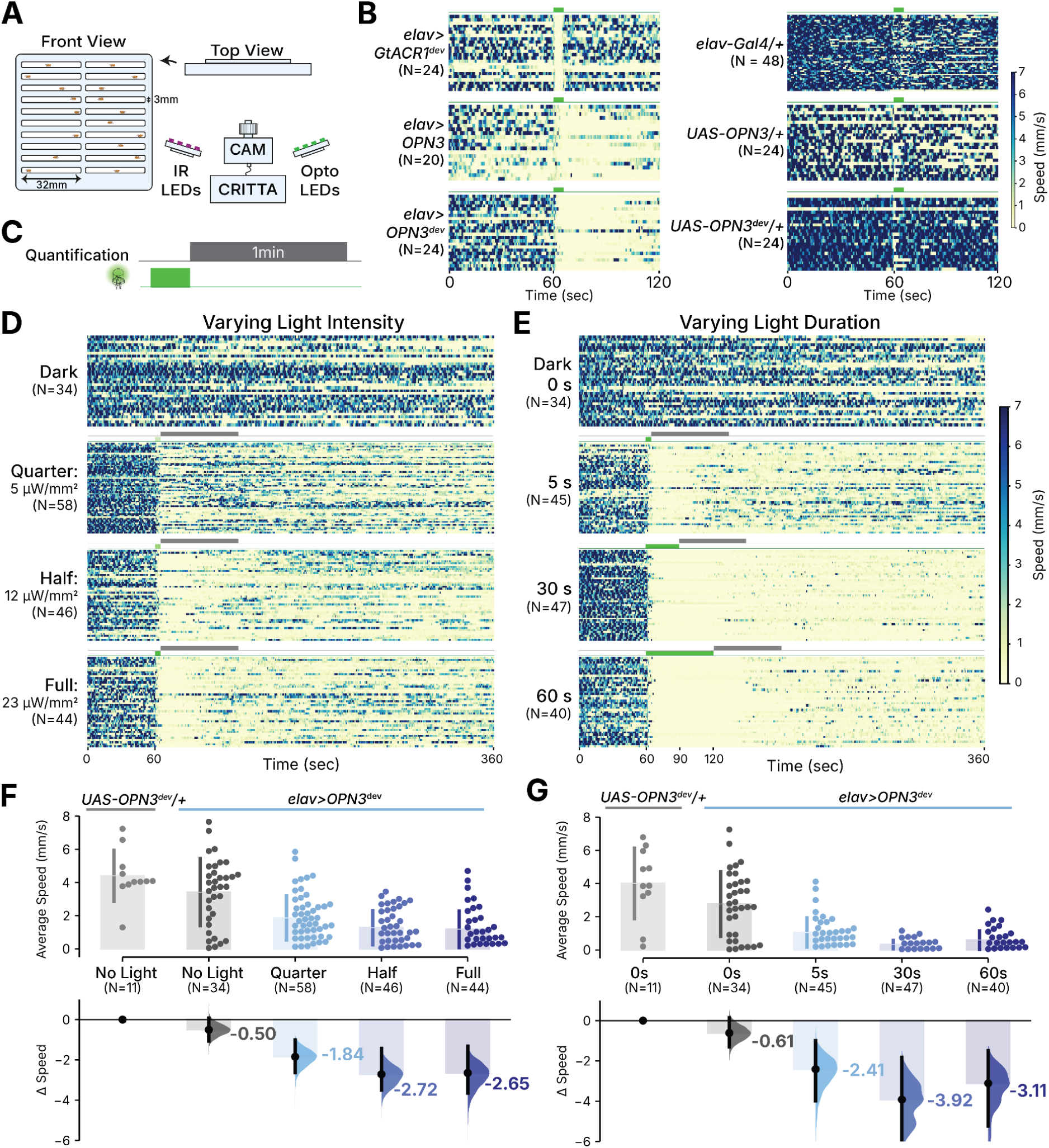
**OPN3 evokes prolonged behavioral inhibition following brief light exposure. (A**) Schematic of the Trumelan apparatus. Flies in individual lanes are tracked from above using IR and CRITTA tracking software. Optogenetic stimulation is delivered via LEDs positioned above the arena. **(B)** Heatmaps of speed over time for *elav-Gal4* crossed to different inhibitory tools: (left) *UAS-GtACR1* (sATR), *UAS-OPN3* (sATR), and *UAS-OPN3*^dev^ and (right) respective driver control (*elav-Gal4/+* on sATR) and responder (*UAS-OPN3/+*) controls raised sATR (middle) and devATR (bottom). Green bars above each heatmap indicate illumination period. **(C)** Inhibition is quantified over a 1 min window (gray bar) immediately following light offset. Light stimulus (green bar) varies in (D) intensity and (E) duration. **(D)** Heatmaps of light intensity (No light, Quarter, Half and Full light intensity) for speed in *elav>OPN3*^dev^ flies. **(E)** Heatmaps of light duration (0 s, 5 s, 30 s, 60 s illumination period) for speed in *elav>OPN3*^dev^ flies. Green bar: illumination period, gray bar: quantification period **(F, G)** Average speed plots 1 min after light offset for *UAS-OPN3*^dev^*/+* controls and *elav>OPN3*^dev^ flies at **(F)** varying light intensities (No Light, Quarter: 4.98 μW/mm^2^; Half: 11.60 μW/mm^2^; Full: 23.48 μW/mm^2^) or **(G)** durations (0 s, 5 s, 30 s, 60 s illumination period), with corresponding Hedge’s g effect sizes on the bottom with 95% confidence intervals. N’s are sample sizes of each group. Superscripted ‘dev’ indicates flies were raised on ATR throughout development and post eclosion.

To determine whether this post-illumination suppression could be titrated, stimulus parameters were manipulated independently and both speed and activity level were quantified over a 1 min window immediately following light offset (**Figure 5C**). Varying light intensity while holding pulse duration constant (5 s) revealed a graded relationship: in darkness, *elav>OPN3^dev^*flies showed only minor differences in baseline locomotion compared to *UAS-OPN3*/+ controls (Δg = −0.50; **Figure 5D, F**), but increasing light intensity reduced post-offset speed up to a plateau, with effect sizes saturating by half-light intensity (Δg = −1.84 for quarter light intensity: 4.98 μW/mm^2^, −2.72 for half light intensity: 11.62 μW/mm^2^, and −2.65 for full intensity: 23.48 μW/mm^2^; **Figure 5F**). Extending illumination duration while holding intensity constant at full light intensity yielded a similar pattern with inhibition saturating by 30s (Δg = −2.41 at 5 s, −3.92 at 30 s, −3.11 at 60 s), with minimal locomotor deficit in the absence of illumination (Δg = −0.61; **Figure 5G**). Activity levels followed a similar pattern across both intensity and duration manipulations (**Supplementary Figure 11**). The point at which inhibition saturated depended on the driver line: *vGAT>OPN3*^dev^ reached maximal suppression with just 5 s of full-light intensity illumination, and locomotion did not resume for approximately 500 s (**Supplementary Figure 12**). Complete effect sizes with 95% CIs for all genotypes across all conditions (genetic controls, light status, food status) are provided in <u>Supplementary Data</u>.

## Discussion

This study evaluated OPN3, a Gα_i/o_-coupled bistable opsin from *Anopheles stephensi*, as an inhibitory optogenetic tool in adult *Drosophila melanogaster* (Mahn et al., 2021). Our goals were to benchmark OPN3 against the established inhibitor GtACR1, and characterize the sustained inhibition produced by GPCR-mediated signaling.

### Limited PdCO efficacy in adult *Drosophila*

There are two relatively straightforward explanations for how PdCO did not inhibit adult *Drosophila* behavior in our hands. First, we used a light intensity comparable to that used for GtACR1, i.e. weaker illumination intensity. PdCO activation in *Drosophila* larvae used 405 nm UV light at 2.7–3.8 mW/mm^2^ ^(Wietek^ ^et^ ^al., 2024)^; sustained PdCO-mediated inhibition by blue light required 2.83 mW/mm^2^ of 470 nm blue light in cuticle-free neuronal cultures (Wietek et al., 2024). Both are approximately 150-fold higher than the 18.3 µW/mm^2^ of 460 nm blue light used in our adult *Drosophila* assays. Wietek and colleagues note that PdCO-mediated inhibition can be short-lived under sub-saturating in vivo light delivery, consistent with our observation (Wietek et al., 2024). Second, PdCO sensitivity peaks near 405 nm (Wietek et al., 2024), and we used LEDs with an emission peak at 460 nm. Worse, the melanized adult cuticle attenuates UV and blue wavelengths far more strongly than the green light (∼530 nm) that activates OPN3 (Kilb, 2021; Malyshev et al., 2017; Scaplen et al., 2021). This cuticular barrier was recognized in *Drosophila* optogenetics and motivated the engineering of red-shifted channelrhodopsins for adult use (Inagaki et al., 2014; Klapoetke et al., 2014). PdCO has been shown to be efficacious in larvae (when actuated with mW of 405-nm light (Wietek et al., 2024)), possibly due to their translucent bodies. In our hands, PdCO did not prove practical for neuronal inhibition in adult flies under conditions comparable to those typically used for ACR1 and OPN3.

### Enhanced membrane trafficking has little effect on OPN3 function

The fly codon-optimized trafficking enhanced variant, eOPN3, did not consistently outperform OPN3. Across *elav-Gal4*, *jus-Gal4*, and *vGAT-GAL4* drivers, the OPN3 construct produced larger effect sizes for both climbing height and speed; only with *nSyb-GAL4* did the trafficking-enhanced variant show substantially stronger inhibition. It has been shown that Kir2.1-derived TS-ER signals can improve opsin membrane expression and photocurrents in cultured mammalian neurons (Gradinaru et al., 2010; Ma et al., 2001; Stockklausner et al., 2001) and enhance axonal targeting in mammalian neurons (Mahn et al., 2021). However, better membrane localization may not always indicate better function. A similar pattern was reported for KCRs in *Drosophila*: KCR1-ET, which carried the same Kir 2.1-derived ER-export and Golgi-trafficking motifs, did not reliably outperform KCR1-GS, with a glycine-serine linker, despite KCR1-ET’s better surface trafficking in Kenyon neurons (Ott et al., 2024). Thus, fusing targeting peptides may not always bestow improved activity upon all opsins for all neuronal types.

### Benchmarking OPN3 against GtACR1

Compared to GtACR1, OPN3 produced comparably robust behavioral inhibition across most neuronal classes tested, including pan-neuronal (*elav-Gal4, nSyb-Gal4*), GABAergic (*vGAT-Gal4*), and glutamatergic (*vGlut-Gal4*) populations. Inhibition was observed with both short and developmentally supplemented ATR, though viability and responses improved with developmental ATR. Under the devATR feeding regime, OPN3 and GtACR1 effect sizes were similar. However, on standard food during development, GtACR1 caused complete developmental lethality in several broad drivers, restricting the conditions under which it can be used. OPN3, on the other hand, was tolerated across all drivers and dietary conditions. This combination of near-equivalent efficacy and improved developmental toxicity positions OPN3 as a practical alternative to GtACR1, particularly for experiments requiring broad neuronal expression.

### Developmental ATR relieves toxicity and enhances opsin efficacy

Unlike mammals and photosynthetic microbes, *Drosophila* raised in lab conditions do not have endogenous ATR, so for channelrhodopsin-based optogenetic experiments, dietary ATR supplementation is required (Dawydow et al., 2014; Nagel et al., 2005; Schroll et al., 2006). However, the timing and duration of ATR supplementation varies across studies; some protocols provide ATR to adult flies for several d prior to experiments (de Vries and Clandinin, 2013; Honjo et al., 2012; Tian et al., 2025; Wu et al., 2014), while others feed ATR starting from the larval stage, albeit at lower concentrations (Aso and Rubin, 2016; Rubin and Aso, 2024; Yamada et al., 2023). The effect of providing ATR earlier in development vs post-eclosion on optogenetic tool efficacy has not been systematically characterized. We found that long-term ATR supplementation during larval development improved both viability and inhibitory efficacy for GtACR1 and OPN3. For GtACR1 specifically, developmental ATR partially rescued the complete lethality observed with broad neuronal drivers. OPN3 showed much lower developmental toxicity compared to GtACR1 regardless of ATR status; whether GPCR opsin apoproteins undergo similar retinal-dependent instability is unknown. One possibility is that the GtACR1 developmental lethality observed under sATR reflects apo-opsin toxicity, paralleling toxic effects in chromophore-deficient mouse photoreceptors (Sakami et al., 2011) and light-independent signaling activity in goldfish cone photoreceptors (Luo et al., 2020). For both GtACR1 and OPN3, developmental ATR feeding produced better inhibition than adult-only feeding, through mechanisms that remain to be determined.

### OPN3 fails to cause paralysis when expressed in OK371 motor neurons

Actuation of OPN3 inhibited behavior across nearly all drivers tested, with one exception: *OK371-Gal4*, which drives expression in many glutamatergic cells, including motor neurons (Hückesfeld et al., 2015; Mahr and Aberle, 2006). This failure was not particular to glutamatergic cells in general, as *vGlut-Gal4*, which labels a larger set of glutamatergic neurons, supported effective behavioral inhibition. We hypothesize that the limitation of OPN3 could be due to a lack of compatible G-proteins or downstream factors, e.g. voltage-gated calcium channels or presynaptic machinery regulated by Gβγ signaling (Mahn et al., 2021). This observation illustrates a general consideration for GPCR-based tools: unlike channelrhodopsins that directly conduct ions, optoGPCRs depend on endogenous signaling components that may vary across cell types. Calcium imaging at OK371 motor neuron terminals would be an informative experiment to address this question.

Duration of post-illumination OPN3 inhibition scales with stimulus parameters Unlike monostable optogenetic inhibitors, where behavioral recovery starts at light offset, OPN3-mediated inhibition persisted for minutes after illumination ended. The magnitude and persistence of the sustained post-illumination suppression scaled with both light intensity and pulse duration up to saturation; this is consistent with an accumulation of stably activated signaling conformers that outlast the photostimulus. In mammalian preparations, eOPN3-mediated synaptic inhibition recovered with a time constant of approximately 5 min (Mahn et al., 2021); we likewise observe sustained, minutes-long behavioral suppression after light offset in adult *Drosophila* with OPN3. The saturation point varied across driver lines: *vGAT-Gal4* reached maximal suppression with just 5 s of illumination (with 23.5 µW/mm^2^ light), while *elav-Gal4* required a longer period of actuation, saturating at 30 s. Additionally, in a memory paradigm, 5 s of illumination using the narrow driver MB320C resulted in strong suppression lasting at least 1 min, though *Th-Gal4* and *MB247-Gal4* required devATR supplementation to achieve the same effect. This driver-dependence suggests that recovery kinetics at least partly reflect properties of the targeted neuronal population: G-protein expression levels, or synaptic architecture, rather than intrinsic characteristics of OPN3. This capacity for sustained inhibition from brief light delivery can reduce the heating (Arias-Gil et al., 2016; Stujenske et al., 2015),phototoxicity (Cardin et al., 2010) and other confounds associated with continuous illumination.

### Limitations

This study has at least four limitations. First, because our readouts are exclusively behavioral, driver-dependent kinetic differences cannot be assigned to cell-autonomous versus network-level causes, and floor (or ceiling) effects on behavior may mask differences in inhibitory strength between conditions. Second, prolonged inhibition is consistent with the molecular bistability described for MosOPN3 (Koyanagi et al., 2013; Mahn et al., 2021), but we cannot exclude alternative or additional contributors such as slow downstream cascade kinetics or network-level effects. Third, OPN3’s failure to inhibit behavior via OK371 cells has an unknown cause; without this knowledge, a user cannot predict which cell types will fall into the OPN3-unresponsive category. Fourth, the improvements in viability and inhibitory efficacy conferred by developmental ATR supplementation are consistent with apo-opsin toxicity, however, we did not test this hypothesis directly. These questions lie beyond the scope of the present study and are suitable subjects for future work.

## Conclusions

Our results verify that OPN3 functions as a robust optogenetic inhibitor; to our knowledge, it is the first inhibitory optoGPCR shown to produce behavioral silencing in adult *Drosophila*, extending OPN3’s utility to a major small-animal model system. For silencing experiments where temporal precision is critical (and the effects of continuous actuating light can be controlled for) channelrhodopsins remain the best choice. With OPN3, a few seconds of green light yields inhibition lasting minutes. So, for experiments examining physiology or behavior over minutes, and/or where actuating light might be confounding, we recommend OPN3. Thus, OPN3 unlocks new experimental capabilities for *Drosophila* inhibitory optogenetics.

## Methods

### *Drosophila* husbandry and all-trans-retinal food

Flies were raised on standard cornmeal-based food containing 1.25% w/v agar, 10.5% w/v dextrose, 10.5% w/v maize and 2.1% w/v yeast at ambient temperature (25°C). All-*trans* retinal (ATR) was supplied as a dietary supplement as it is in spontaneous thermal equilibrium with 13-*cis* retinal, which OPN3 primarily binds to form a functional photopigment (Koyanagi et al., 2013). ATR (Sigma-Aldrich, R2500) was dissolved in 100% ethanol in the dark and mixed with warm liquefied food to a final standard concentration of 0.5 mM. All ATR food vials were kept at 4°C and used within 2 weeks of preparation. The short ATR (‘sATR’) protocol consisted of raising flies on standard food without ATR through development and eclosion, then transferring them to ATR-supplemented food for 48–72 h immediately prior to behavioral experiments. Developmental ATR supplementation (‘devATR’) consisted of crossing parental flies on ATR food and raising offspring on ATR food continuously through parental mating, embryogenesis, larval growth, eclosion and adulthood—between 14 to 17 d. All flies used for behavioral experiments were 5–7 d post-eclosion at the time of testing, irrespective of ATR feeding regime.

### Drosophila stocks

The following stocks were obtained from the Bloomington Drosophila Stock Center (BDSC):

*OK371-Gal4* (BDSC#26160), *elav-Gal4* (BDSC #458), *nSyb-Gal4* (BDSC #51635), *vGAT-Gal4* (BDSC #58980), *vGLUT-Gal4* (BDSC #24635), *jus-Gal4* (BDSC #46416; otherwise known as *GMR59D01-GAL4*), *Th-Gal4* (BDSC #8848; otherwise known as *ple-Gal4*), *MB247-Gal4* (BDSC #50742; otherwise known as *Mef2-Gal4.247*), MB320C (BDSC #68253). The *UAS-ACR1* stock was generated as described in (Mohammad et al., 2017). The *UAS-PdCO* line was graciously donated by Dr Peter Soba.

### Fly constructs and genetics

Two opsin transgenes were built for this study, UAS-OPN3 and UAS-eOPN3, both based on the sequence of pAAV-CKIIa-eOPN3-mScarlet-WPRE (Addgene #125712; (Mahn et al., 2021))), which carried the 99 aa C-terminally truncated MosOPN3 cDNA (residues 1–330, the MosOpn3ΔC variant (Koyanagi et al., 2013); GenBank AB753162). For UAS-OPN3, an alanine triplet (AAA) linker and eYFP reporter were fused in-frame at the C-terminus, with the Rho1D4 affinity-tag epitope (TETSQVAPA) carried over from the original design (Koyanagi et al., 2013). For UAS-eOPN3, the cassette uses three hemagglutinin peptides, 3×HA (3×YPYDVPDYA), as the epitope tag and mScarlet3 (Gadella et al., 2023) fluorophore. Two Kir2.1-derived trafficking elements were incorporated following the KCR-ET design (Ott et al., 2024): a 20-residue trafficking signal KSRITSEGEYIPLDQIDINV (Hofherr et al., 2005) sits at the C-terminus of OPN3 immediately upstream of the 3×HA tag, and a 7-residue ER export motif (FCYENEV; (Ma et al., 2001; Stockklausner et al., 2001) caps the cassette downstream of mScarlet3. Visualization of inserts can be found in Supplementary Figure 1A. Both opsin coding sequences were codon-optimized for Drosophila and synthesized by Genscript de novo. The OPN3 cassette was inserted into pJFRC7-20×UAS-IVS-mCD8::GFP (Addgene #26220) at the XhoI/XbaI sites, downstream of the hsp70 basal promoter and 20×UAS array, in place of the original mCD8::GFP insert. The eOPN3 cassette was cloned into pJFRC81-10×UAS-IVS-Syn21-GFP-p10 (Addgene #36432) at the XhoI/XbaI sites, downstream of the hsp70 basal promoter and 10×UAS array, in place of the parental Syn21-GFP-p10 insert. The two synthesized cassettes were embryo-injected by BestGene for ϕC31 integration into attP2 (chromosome 3), and resulting transgenics were balanced over TM6C and maintained as homozygous stocks.

Expression was checked under confocal imaging using the eYFP (OPN3) or mScarlet3 (eOPN3) reporter signal. For all behavioural assays, F1 progeny of a driver-Gal4 > UAS-responder cross served as experimental subjects while driver-Gal4 and UAS-responder flies were each crossed with w^1118^ flies each and the F1 progeny of those were then used as controls.

### Immunohistochemistry and confocal imaging

Adult brains were dissected in cold phosphate-buffered saline (PBS; 0.1 mM PB) and fixed with 4% paraformaldehyde for 20 min as previously described (Mohammad et al., 2017). Fixed brains were washed 3 × 15 min in PBST and incubated in primary antibodies in PBST at 4°C for 48 h, after which they were rinsed and incubated in secondary antibodies in PBST at 4°C for 24 h. Finally, the brains were washed 3 × 15 min in PBS mounted on microscope slides in Vectashield (Vector Laboratories) and covered with a coverslip. Primary antibodies used were 1:50 mouse anti-DLG (Developmental Studies Hybridoma Bank; RRID AB_528203) and 1:1000 chicken anti-GFP (Abcam; RRID ab13970). Secondary antibodies used were 1:200 Alexa Fluor goat anti-mouse 568 (Thermo Fisher; RRID AB_2534072) and 1:200 Alexa Fluor goat anti-chicken 488 (Thermo Fisher; RRID AB_2534096). Slides containing mounted fly brains were imaged on a Zeiss LSM700 upright microscope using a 20X objective. Pixel intensities were summed across z-stack slices in ImageJ (Sum Slices Z-projection).

### Opsin expression in N2a cell culture

The OPN3 construct was cloned into the multiple cloning sites of a pcDNA3.4 vector (Genscript). 500ng of the respective DNA constructs were transfected into N2a cells (ATCC, CCL-131™) using Lipofectamine 3000 (Invitrogen). The cells were left to incubate in serum-free media for 48 h. The N2a cultures were then washed three times with PBS and fixed for 20 min at room temperature with 4% paraformaldehyde diluted in PBS-Triton X-100 (0.25%, Thermo Fisher Scientific 85111). After fixation, the cells were blocked in 5% BSA (Gold Biotechnology, A-420-500) diluted in PBS-Triton X-100 (0.25%) for 1 h at room temperature and stained for GFP (Abcam ab13970, RRID: AB_300798) at 1:2000 v/v dilution for 1.5 h at 37°C. Afterward, the cultures were rinsed three times with PBS and incubated with an Alexa 488 goat anti-chicken (Thermo Fisher Scientific A-11039, RRID: AB_2534096) at 1:500 v/v dilution and DAPI at 1:1000 v/v dilution for 1 h at 37°C. Finally, the cells were washed three times with PBS and mounted onto microscope slides in Vectashield Vibrance mounting media (Vector Laboratories, H-1700). Imaging was performed on a Zeiss LSM700 upright microscope using a 100× objective.

### Optogenetic illuminations

Green (peak emission 530 nm, Luxeon Rebel, SP-05-G4, Quadica Developments Inc.) and blue LEDs (peak emission 460 nm, Luxeon Rebel, SP-05-B4, Quadica Developments Inc.) were mounted on heatsinks (Luxeon Star N50-25B, Quadica Developments Inc.) for all optogenetic experiments. LEDs were powered by the output voltage of a 700-mA BuckPuck driver (Luxeon Star 3023-D-E-700, Quadica Developments Inc.). The experimental setup, including the illumination system and behavioral apparatus, was housed inside a temperature-regulated incubator (MIR-154, Sanyo) for the duration of the optogenetic experiments. Light-intensity measurements were performed using a photodiode (Thorlabs S130C) connected to a power and energy-meter console (Thorlabs PM100D) in a dark room, and zeroed before each measurement.

### Viability assay

To assess for developmental toxicity, eight virgin females, aged for 5 to 10 d, were crossed with four males per vial. After two days, the flies were transferred to a population cage containing a fresh grape-juice agar plate and incubated at 25°C under a light/dark cycle.

Grape-juice plates were prepared by combining 1% agarose (Bio-Rad, 1613102), grape concentrate (Welch’s), yeast solution (MP Biomedicals, 0290331225), and nipagen (Megachem Limited), then poured into 6 cm cell culture plates. After 24 h, the plate was collected and replaced with a fresh agar plate; this was repeated to obtain three replicate plates per experimental condition. Embryos were counted manually and then transferred to a 6 cm cell culture plate containing regular fly food and sealed within a larger 15 cm plate. Plates were incubated until eclosion (∼10–15 d). Flies were then anesthetized, counted, and any unhatched embryos recorded. Lethality proportion was calculated per genotype as the fraction of hatched embryos that failed to eclose as adults (1 − eclosed/hatched), pooled across replicate vials, and the paired mean difference between the hatching baseline and eclosion endpoint was calculated using Dabest (Lu et al., 2026).

### Climbing assay

*Drosophila* climbing behavior was measured in a 9 mm-thick 170 × 94 mm acrylic cassette. Seventeen chambers were cut into a 3 mm-thick transparent acrylic sheet and fixed onto a white acrylic diffuser piece (See Figure 3A for schematic). After ice anesthesia of 30 s, individual flies were transferred into a single rectangular climbing chamber (7W × 86H × 3D mm). The flies were given 3 min to recover from anesthesia before the start of the experiment. Climbing behavior was recorded at 5 fps under infrared backlighting (850 nm). Recording was performed with a Chameleon3 near-infrared video camera (FLIR CM3-U3-13S2C) equipped with a 4.4–11 mm FL High Resolution Varifocal Lens (Edmund Optics) and a 850 nm longpass filter (Green.L, 58-850). The cassette was illuminated from the front with 7 LEDs at a distance of 26cm. Each video frame was processed in real time connected to CRITTA tracking software (Krishnan et al., 2014). The cassette was first manually tapped downwards before the tracking started and flies were then allowed to climb for 23 s in the dark (infrared light only). The tracking system was briefly paused while the flies were re-agitated. After agitation, the flies were given 3 s to climb in the dark, after which the LED lights were activated for 20 s (green at 22 μW/mm^2^ or blue light at 18.3 μW/mm^2^). The LED lights were then turned off, and flies were recorded in the dark for additional 20 s before the experiment was concluded. Climbing height was calculated per fly as the mean vertical position (mm above chamber floor) during each light-on, light-off epoch. Speed was calculated per fly as the mean instantaneous speed (mm/s) over the same epoch, computed from frame-to-frame centroid displacement.

### Associative olfactory learning assay

For associative olfactory learning, we used an in-house developed assay called the Multi-fly Olfactory Trainer (MOT; see Figure 4B for schematic) (Kan et al., 2021; Mohammad et al., 2017; Ott et al., 2024), which consists of 10 rectangular chambers of dimensions 50 × 5 × 1.3 mm cut into white acrylic. To facilitate shock delivery, the floor and ceiling of each chamber were made from glass with indium tin oxide (ITO) electrodes printed onto the surface (Visiontek UK). Behavior was recorded at 20 frames per second (fps) using a Guppy F-080 camera (Allied Vision Technologies) connected to a PCI-1409 (National Instruments) video acquisition board. Carrier air flow rate was 500 mL/min; the odors used were 9 parts per million (ppm) 4-methylcyclohexanol (MCH) and 6 ppm 3-octanol (OCT). For optogenetic actuation, green LEDs with a peak wavelength of 525 nm and intensity of 19.9 μW/mm^2^ (Luxeon Star).

The typical conditioning protocol used in our experiments is shown in Figure 4A. Flies were first agitated using air puffs, then presented with both odors for a 2-min odor-bias test, followed by 1 min training with shock against the conditioned stimulus (CS+) odor, 1 min training of the non-CS odor (CS-) with no shock, a 2-min static test where both odors are presented with no agitation, and finally a 2-min agitation test where flies are agitated with airpuffs prior to odor presentation. For optogenetic actuation, flies were either presented with constant light during the shock epoch (Figure 4C-E, Supplementary Figure 8), or a short burst of light ranging from 1-10 s directly prior to the shock (Figure 4G-H, Supplementary Figures 9-10). 5–7 flies were run per chamber. The shock protocol used was 60 V shock, 1 s per shock, delivered 12 times over 1 min. The last 30 s of the bias test and agitation test are used for analysis; only the agitation test is used because flies often only express weak or no learning during the static test, and agitation is required to induce expression of learning (data not shown; manuscript in preparation). A preference index is calculated as follows for each of the bias and agitation test epochs:

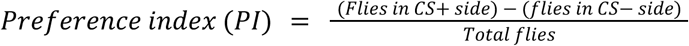

A ΔPI is then calculated between bias and agitation test, and a ΔΔPI calculated between experimental flies and genetic control groups’ ΔPI values (defined as driver-only or responder-only flies):

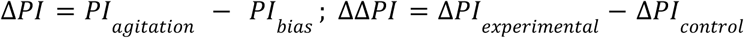

For experiments involving shorter periods of light stimulation (for 1 s, 3 s, 5 s or 10 s), light was presented directly prior to the shock epoch.

% learning compared to controls is calculated as such:

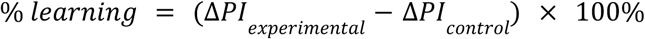

By performing this calculation across the full bootstrap distribution for both ΔPIs, we are able to obtain a bootstrapped distribution of % learning, and the bootstrapped distribution along with means and 95% confidence intervals are then plotted.

### Trumelan activity-monitoring assay

The Trumelan cassette comprised a grid of 26 individual fly chambers arranged in two columns of 13 chambers, each measuring 32 × 3 × 3 mm; width × height × depth).

Chambers were milled into a 3-mm-thick acrylic plate, sealed with a backing acrylic sheet, and enclosed on the front by an additional acrylic cover (See Figure 5A for schematic). To enhance visual contrast, a sheet of matte black card was positioned behind the cassette. The assembled cassette was mounted vertically inside a temperature-controlled incubator (Sanyo MIR-154) maintained at 25°C. Behavioral recordings were acquired at 10 frames/s using a near-infrared FLIR Grasshopper3 camera (Edmund Optics, GS3-U3-41C6NIR-C) fitted with a 50 mm fixed-focus lens (Edmund Optics, VS-C5024-10M) and an 850 nm long-pass filter (Green.L, 58-850). Continuous illumination was provided by two arrays of infrared LEDs with a peak emission at 850 nm. Video streams were analyzed online using CRITTA LabVIEW software to extract behavioral parameters for each fly. Each experiment consisted of three sequential phases: an initial 60 s dark baseline (infrared illumination only), followed by exposure to green LED light (23.48 µW/mm^2^), and a final 60 min dark period. Both the intensity (0, 4.98, 11.60, or 23.48 µW/mm^2^; referred to as No Light, Quarter, Half, and Full) and duration of the green LED phase (0 s, 5 s, 30 s, 60 s) varied across experiments and are specified in the corresponding figure legends.

## Statistical analyses

Experiments were not performed in a blinded fashion. No initial power calculations were made to determine the sample sizes; significance tests were not conducted (Cumming and Calin-Jageman, 2017). Estimation statistics were used to analyze quantitative data with the DABEST software library (Ho et al., 2019; Lu et al., 2026). Hedges’ *g*, a standardized effect size representing the mean difference divided by the pooled standard deviation, was computed using the pandas, scikits-bootstrap, seaborn, and SciPy packages. Data analysis was performed and visualized with Jupyter Python notebooks using DABEST.

## Data Availability

The data and code that support the findings of this study can be found on Zenodo repositories (DOIs: 10.5281/zenodo.20323114 and 10.5281/zenodo.20347628).

## Acknowledgements

We are grateful to James Stewart and Joses Ho for assistance with the Climbing, Trumelan and OSAR rig. We thank Dr Peter Soba (Friedrich-Alexander University Erlangen-Nürnberg) for donating the PdCO stock. We thank Dr Jun Nishiyama (Duke-NUS Medical School) for sharing the N2a mouse neuroblastoma cells.

## Author Contributions

Conceptualization: NL and ACC; Experiment design: NL, YM and ACC; Methodology: NL, YM; Software: NL, YM; Data Analysis: NL, YM; Investigation: NL (construct design, climbing, fecundity and Trumelan experiments, data analysis, data processing), YM (Immunohistochemistry, MOT experiments, data analysis, data processing), LD (Climbing, Trumelan and fecundity experiments), DS (Cell transfection, immunohistochemistry, microscopy), JA (Trumelan experiments)

## Resources

Writing – Original Draft: NL; Writing – Revision: NL, YM and ACC; Visualization: NL, YM and ACC; Supervision: NL and ACC; Project Administration: ACC; Funding Acquisition: ACC.

## Supplementary Figures

**Supplementary Figure 1.**
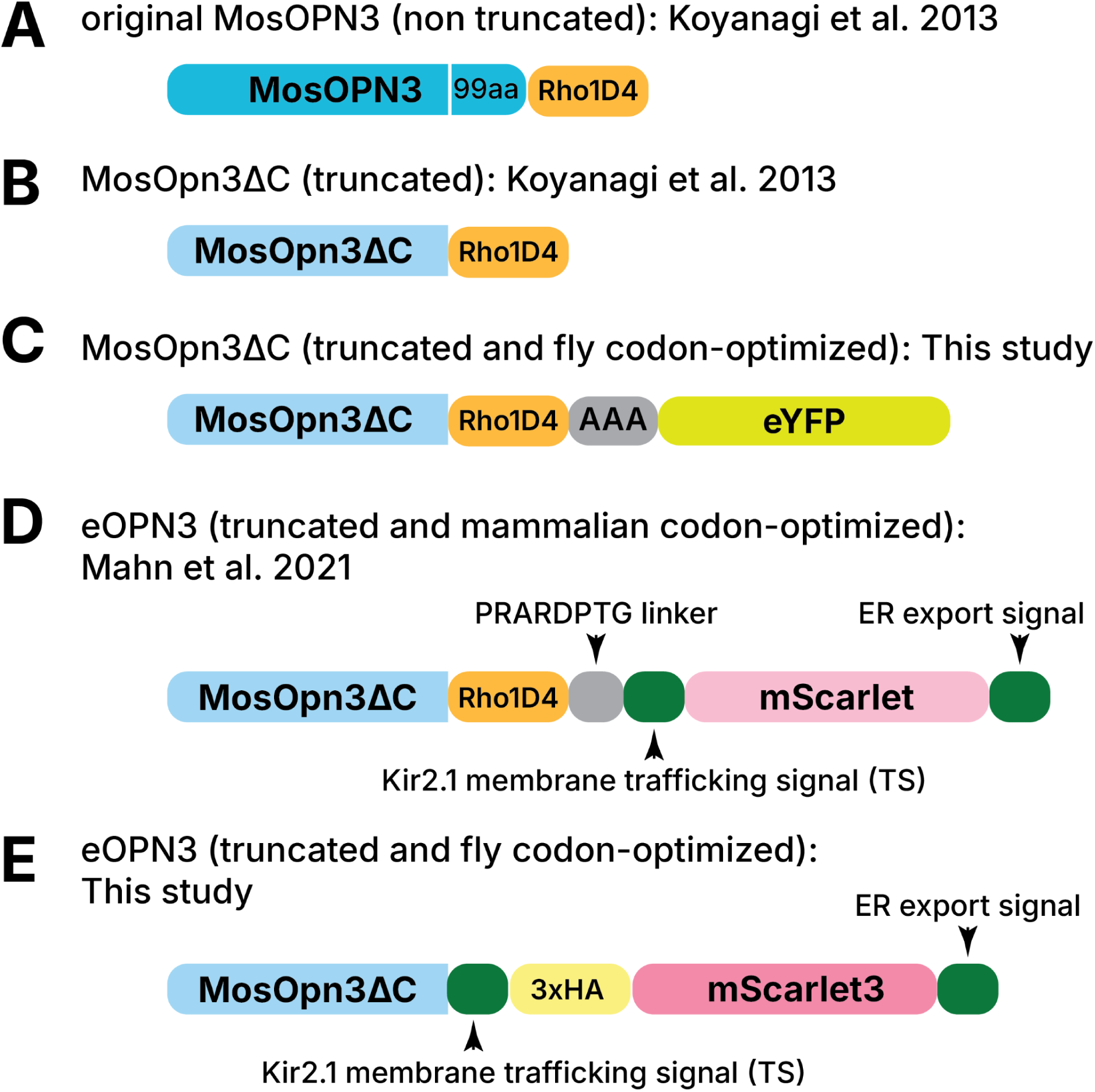
Summary of OPN3-based constructs. (**A**) Full-length MosOPN3 (dark blue) and Rho1D4 epitope tag (orange) (Koyanagi et al., 2013) (**B**) 99aa C-terminal truncated MosOPN3 variant (MosOPN3ΔC, light blue; residues 1–330) with Rho1D4 (Koyanagi et al., 2013). (**C**) Fly codon-optimized version of (B) with AAA linker (gray) and eYFP reporter (yellow-green). Same construct shown in Fig. 1A, reproduced here for clarity. (**D**) Mammalian eOPN3 with PRARDPTG linker (grey), Kir2.1 membrane trafficking signal (green), mScarlet reporter (light pink), and Kir2.1 ER export motif (green) (Mahn et al., 2021) (**E**) Fly codon-optimized eOPN3 of this study, with Kir2.1 membrane trafficking signal, 3×HA epitope tag (yellow), mScarlet3 reporter (pink), and the Kir2.1 ER export motif.

**Supplementary Figure 2.**
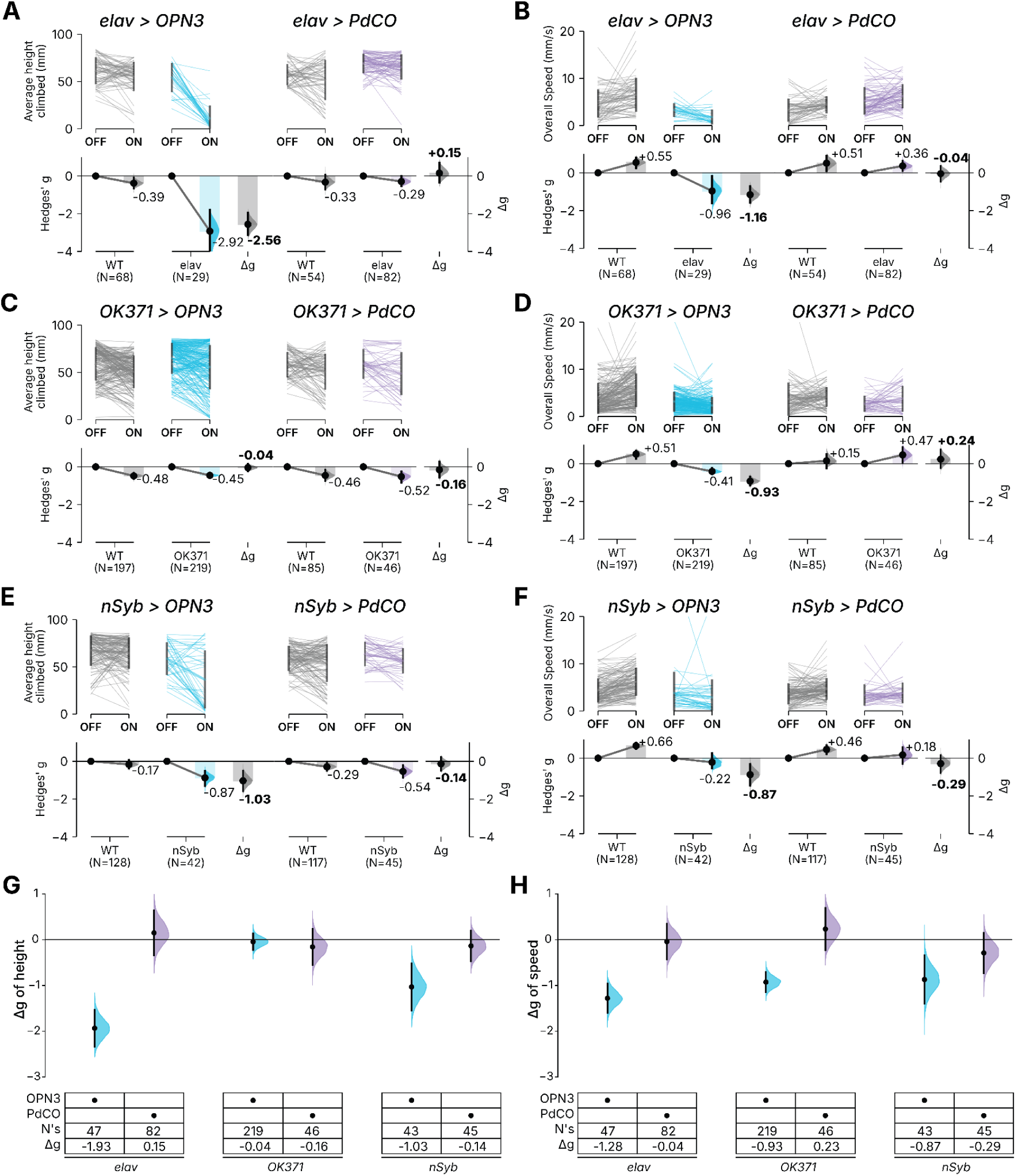
PdCO shows limited efficacy in adult *Drosophila* compared to OPN3. (**A–F**) Individual Δ*g* height (**A**, **C**, **E**) and speed (**B**, **D**, **F**) estimation plots comparing light OFF and ON in wild type and controls flies expressing (**A**, **B**) *elav-Gal4*, (**C**, **D**) *OK371-Gal4*, and (**E**, **F**) *nSyb-Gal4* driving PdCO and OPN3 (sATR). (**G, H**) Summary of Δ*g* height (**G**) and speed (**H**) effect sizes across the three drivers; gridkey indicates driver and opsin, sample size (N’s), and Δg effect size. Purple, PdCO; blue, OPN3. Error bars, 95% CI. Note that the *elav>OPN3* and *nSyb>OPN3* flies are the same animals shown in Figure 3E; the height data are redrawn from Figure 3E, while the speed data are an additional metric computed from the same flies, both included to enable direct graphical comparison.

**Supplementary Figure 3.**
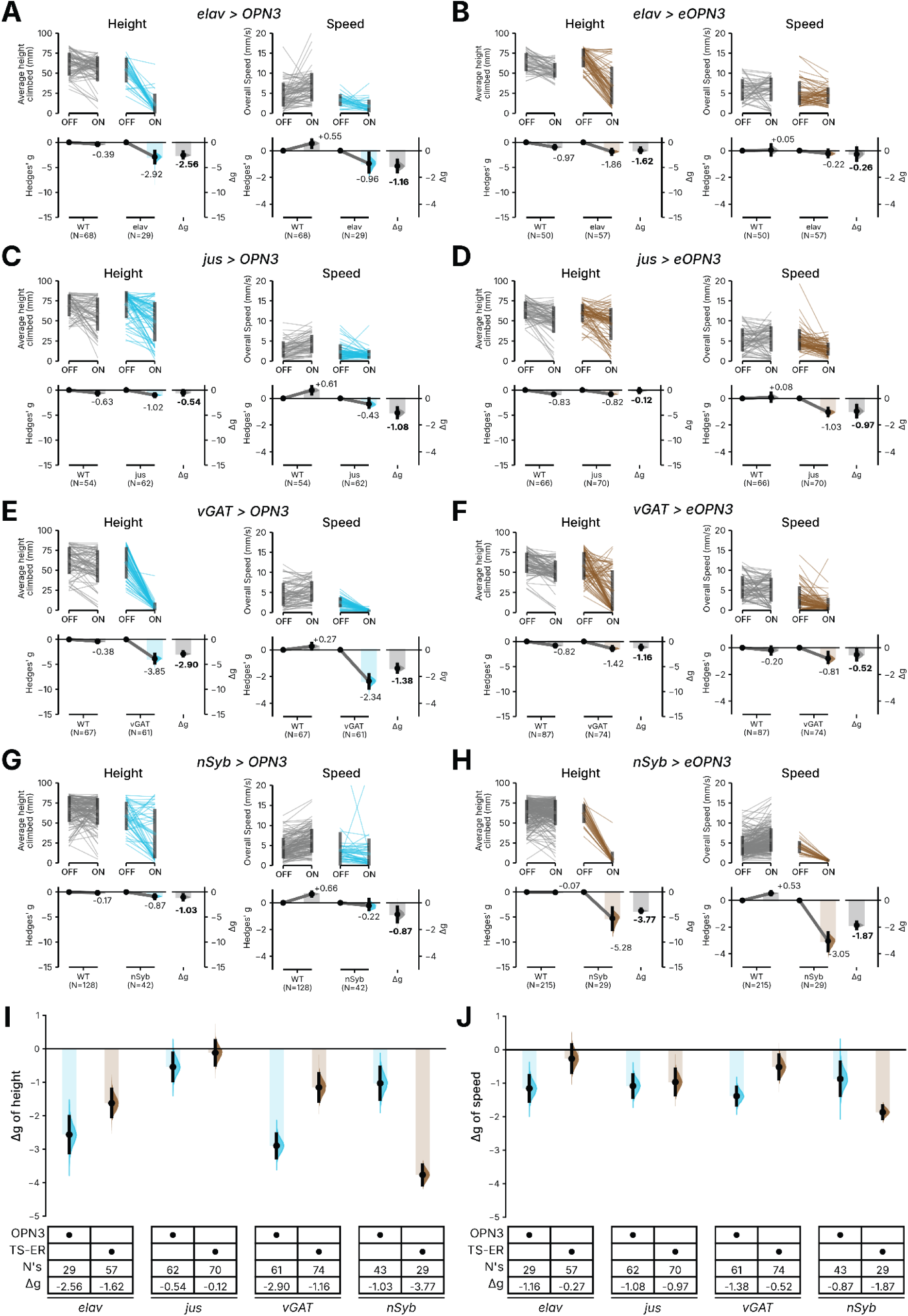
Addition of a *Kir2.1* trafficking signal reduces eOPN3 efficacy in most drivers. (**A–H**) Individual Δ*g* height (left subpanels) and speed (right subpanels) estimation plots comparing light OFF and ON in wild type and controls flies expressing (**A**) *elav-Gal4>OPN3*, (**B**) *elav-Gal4>eOPN3*, (**C**) *jus-Gal4>OPN3*, (**D**) *jus-Gal4>eOPN3*, (**E**) *vGAT-Gal4>OPN3*, (**F**) *vGAT-Gal4>eOPN3*, (**G**) *nSyb-Gal4>OPN3*, and (**H**) *nSyb-Gal4>eOPN3* (**I, J**) Summary of Δ*g* height (**I**) and speed (**J**) effect sizes across the four drivers; gridkey indicates driver and opsin, sample size (N’s), trafficking signal sequence and Δ*g*. Error bars, 95% CI. Note that the *elav>OPN3*, *jus>OPN3*, *vGAT>OPN3* and *nSyb>OPN3* flies are the same flies shown in Figure 3E. Height data for all four are redrawn from Figure 3E. Speed data for *elav>OPN3* and *nSyb>OPN3* are redrawn from Supplementary Figure 2, while speed data for *jus>OPN3* and *vGAT>OPN3* are an additional metric first computed here from the same flies in Figure 3E. All are included to enable direct graphical comparison with eOPN3.

**Supplementary Figure 4.**
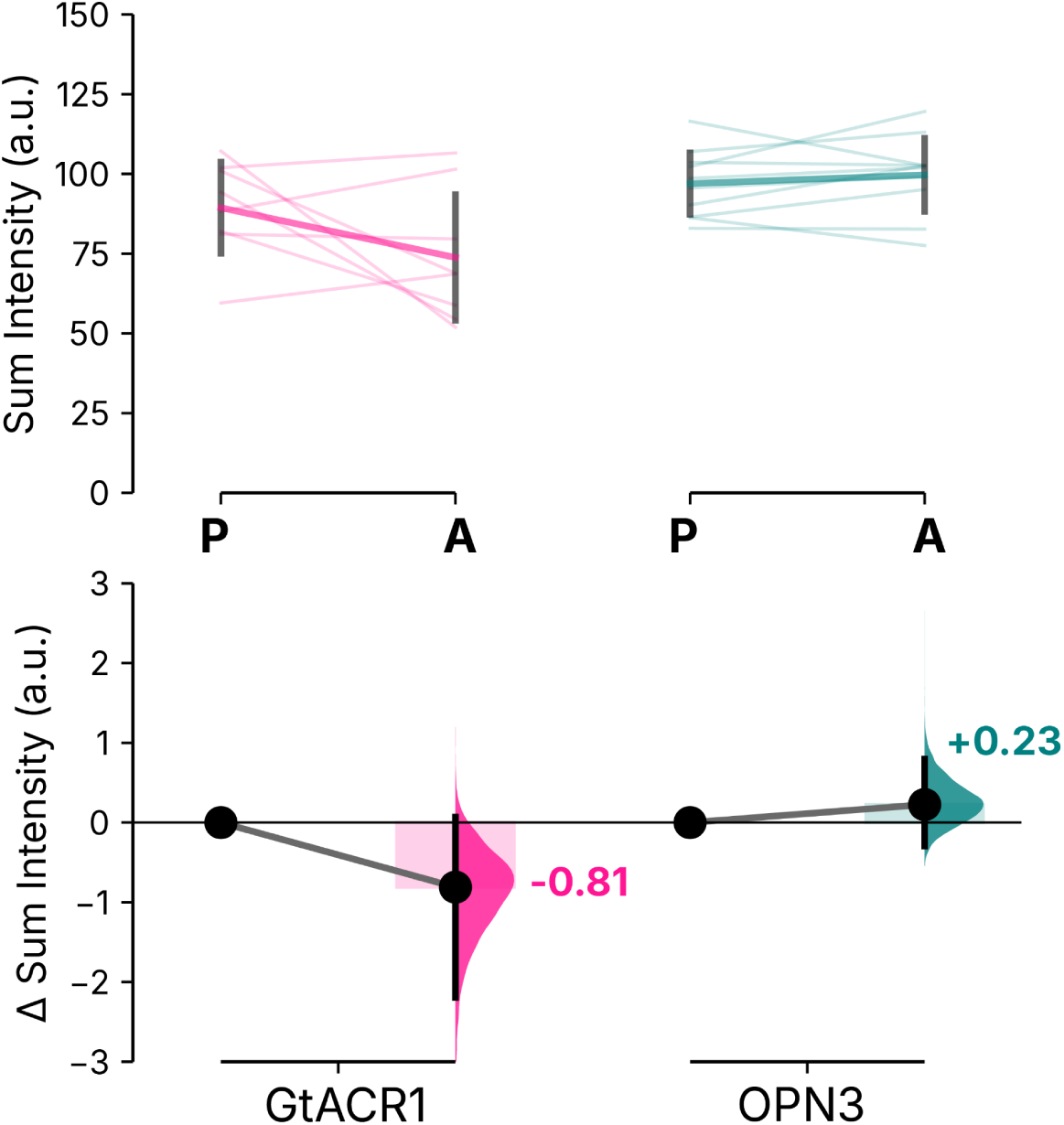
Quantification of brain regions of MB247 for GtACR1 and OPN3. Quantification of anti-GFP pixel intensities summed across z-stack slices in posterior (P) versus anterior (A) brain regions for each opsin transgene crossed with *MB247-Gal4*. Top: Individual brain regions shown as slope plots. Height of the bars shows average intensity values. Bottom: posterior - anterior Hedges’ *g* of anti-GFP intensities. Error bars represent 95% CI. n = 8 hemispheres per genotype

**Supplementary Figure 5.**
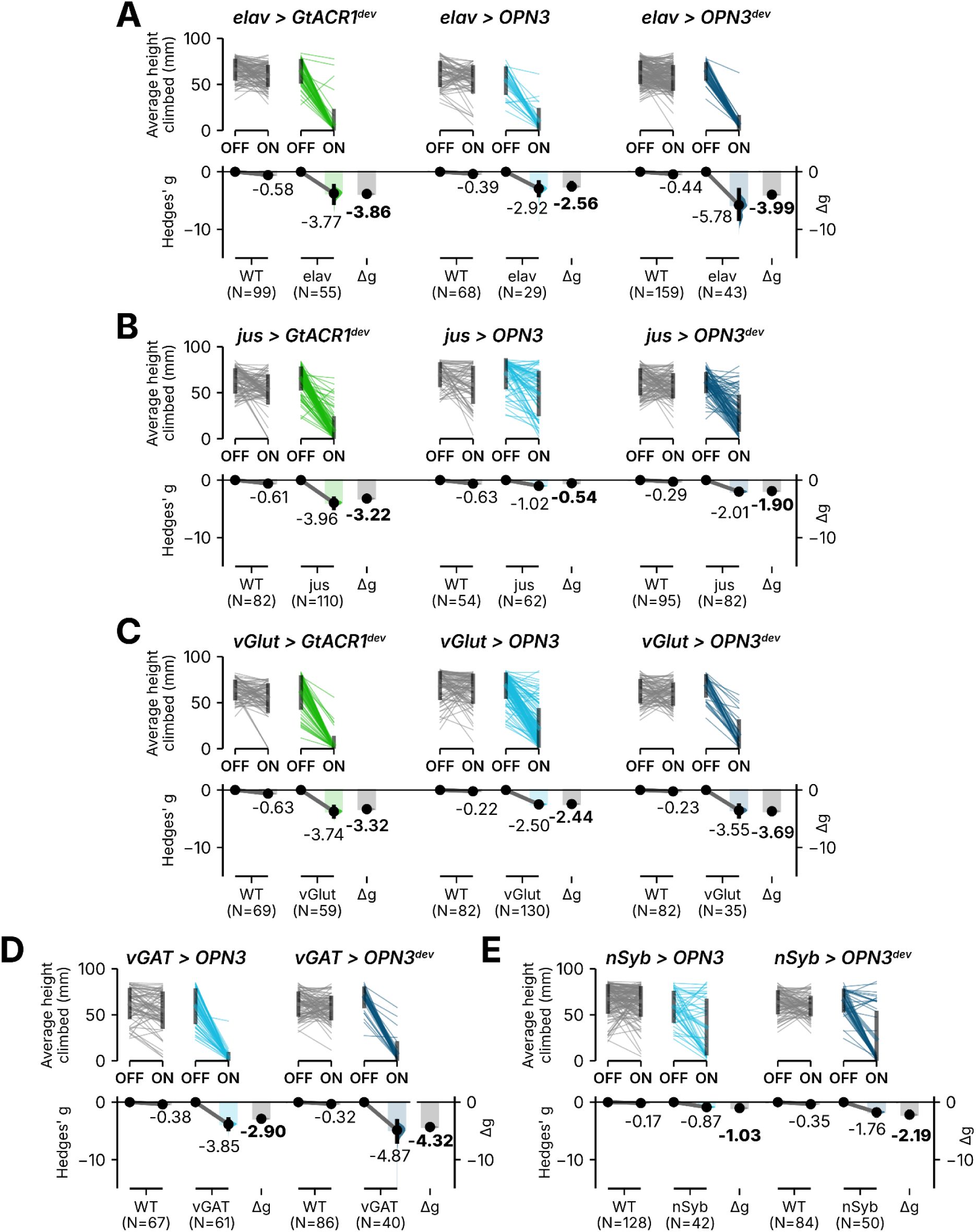
Individual and summary plots of Δg height effect sizes across multiple neuronal Gal4 drivers. Individual Δg height estimation plots comparing light OFF and ON in wild type and controls flies expressing (**A**) *elav-Gal4*, (**B**) *jus-Gal4*, (**C**) *vGlut-Gal4* driving opsins GtACR1, OPN3, and OPN3^dev^ (**D**) *vGAT-Gal4* and (**E**) *nSyb-Gal4* driving OPN3 and OPN3^dev^; Green, GtACR1; blue, OPN3; darker shades, devATR; lighter shades, sATR. Error bars, 95% CI. Superscripted ‘dev’ indicates flies were raised on ATR throughout development and post eclosion. Note that the *elav*, *jus*, *vGlut*, *vGAT* and *nSyb* data for GtACR1^dev^, OPN3 and OPN3^dev^ are the same flies and height measurements shown in the Figure 3E summary plot, presented here as individual estimation plots.

**Supplementary Figure 6.**
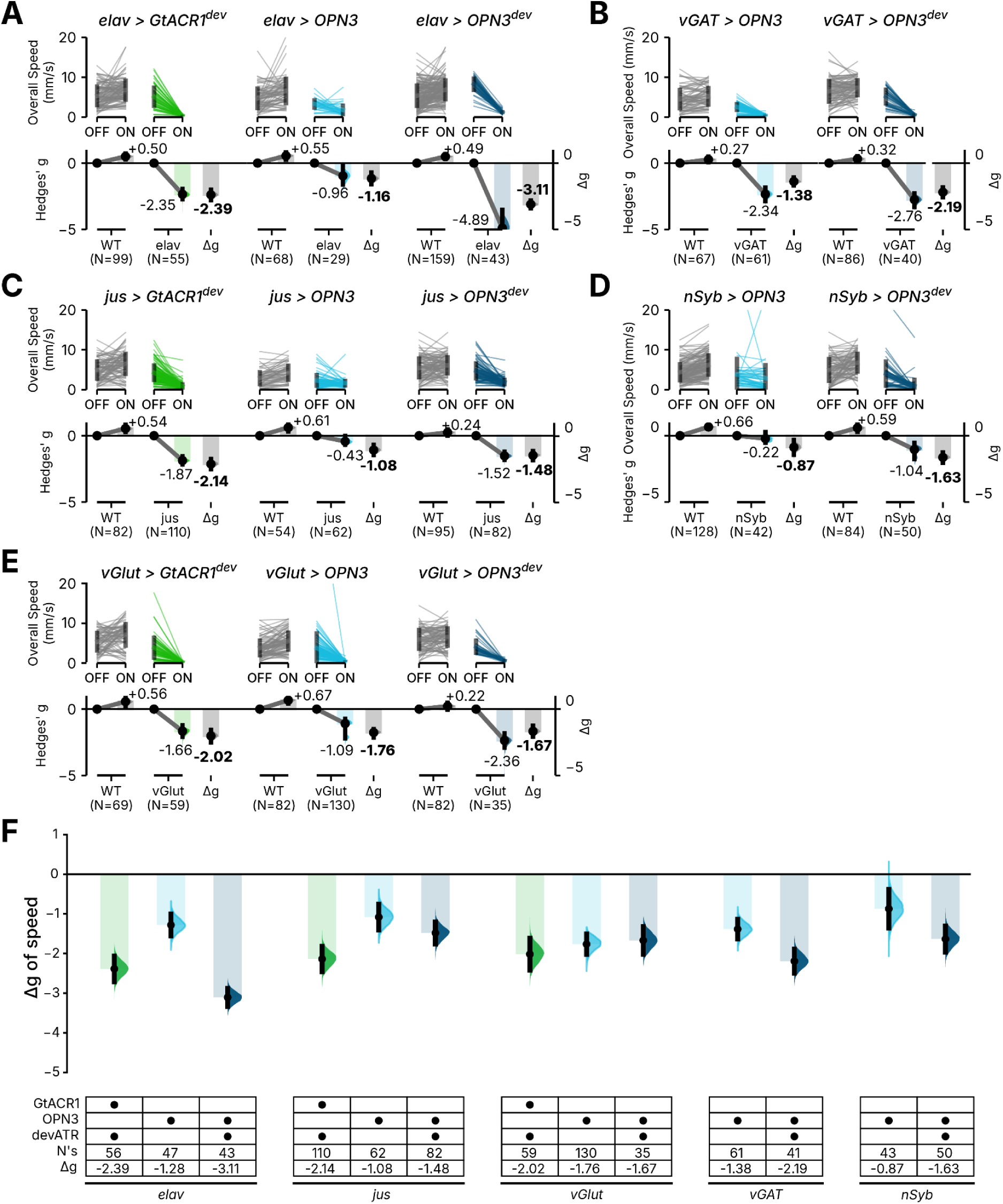
Individual and summary plots of Δg speed effect sizes across multiple neuronal Gal4 drivers. Individual Δg speed estimation plots comparing light OFF and ON in wild type and controls flies expressing (**A**) *elav-Gal4*, (**C**) *jus-Gal4*, (**E**) *vGlut-Gal4* driving opsins GtACR1, OPN3, and OPN3^dev^ (**B**) *vGAT-Gal4* and (**D**) *nSyb-Gal4* driving OPN3 and OPN3^dev^; Green, GtACR1; blue, OPN3; darker shades, devATR; lighter shades, sATR. Error bars, 95% CI. (**F**) Summary of Δ*g* effect sizes across all drivers, opsins, and ATR conditions; gridkey lists drivers, opsins, ATR status, sample size (N), and Δ*g*. Green, GtACR1; blue, OPN3; darker shades, devATR; lighter shades, sATR. Error bars, 95% CI. Superscripted ‘dev’ indicates flies were raised on ATR throughout development and post eclosion. Note that the *elav*, *jus*, *vGlut*, *vGAT* and *nSyb* data for GtACR1^dev^, OPN3 and OPN3^dev^ are the same flies shown in Figure 3E summary plot; speed is an additional metric to the height presented there. Speed for *elav>OPN3* and *nSyb>OPN3* is redrawn from Supplementary Figure 2, *jus>OPN3* and *vGAT>OPN3* from Supplementary Figure 3, with the remaining speed data computed here from the same flies. All are presented as individual estimation plots to enable direct graphical comparison.

**Supplementary Figure 7.**
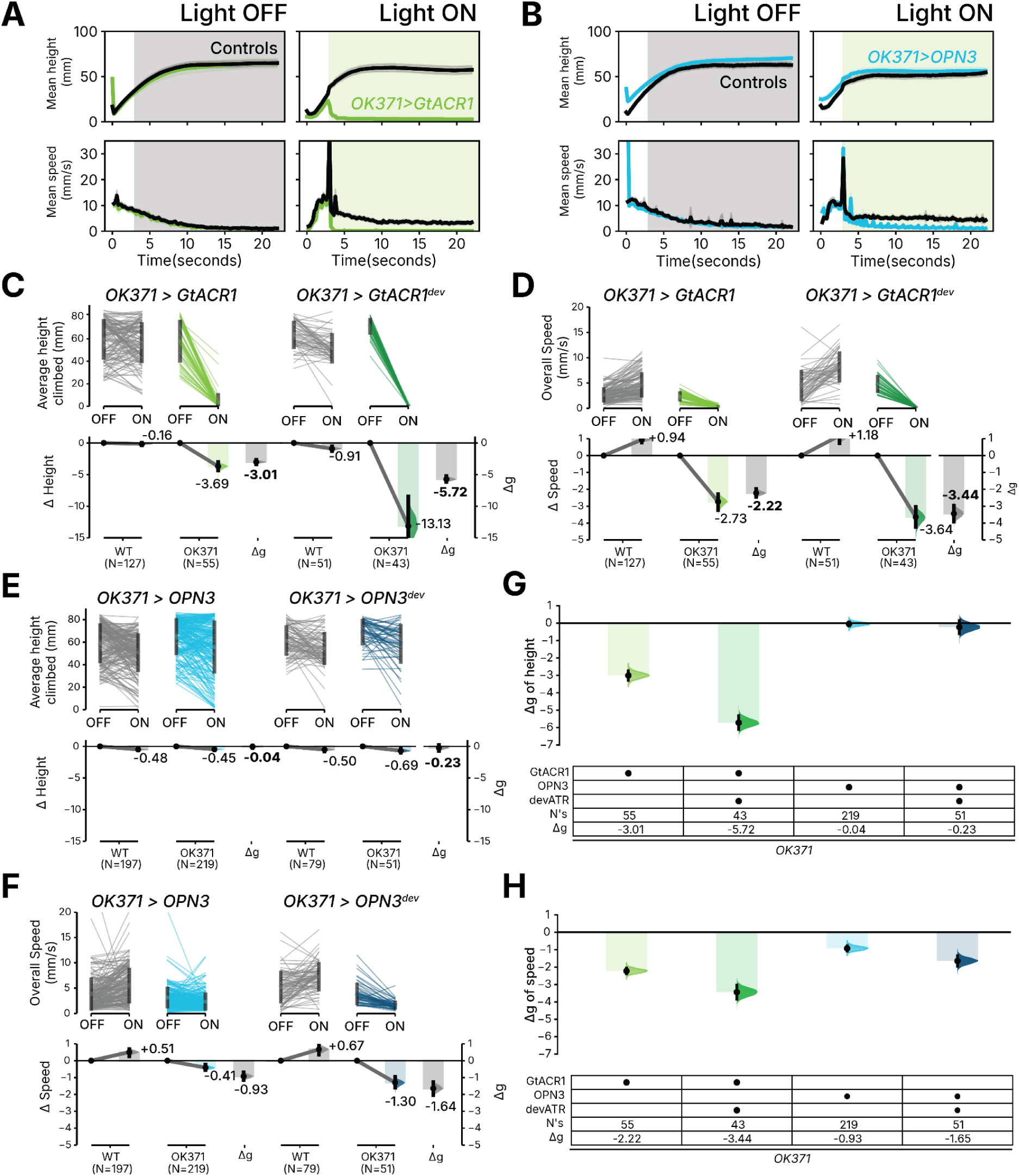
*OK371-GAL4* glutamatergic neurons are not inhibited by OPN3. Time-series traces showing mean climbing height (top) and speed (bottom) for genetic controls and experimental crosses expressing (**A**) *OK371>GtACR1* (n=55) and (**B**) *OK371>OPN3* (n = 219). Shaded regions denote light off or on in the experiment sequence; shaded regions of line plots represent SEM. (**C**, **D**) Individual Δ*g* plots for *OK371-Gal4* driving GtACR1 and GtACR1^dev^ for climbing height (**C**) and speed (**D**). (**E**, **F**) Individual Δ*g* plots for *OK371-Gal4* driving OPN3 and OPN3^dev^ for climbing height (**E**) and speed (**F**). Superscripted ‘dev’ indicates flies were raised on ATR throughout development and post eclosion. (**G**, **H**) Summary of Δ*g* effect sizes across opsins and ATR dietary conditions for height (**G**) and speed (**H**); gridkey indicates driver and opsin, ATR status, sample size (*N*), and Δ*g*. Green, GtACR1; blue, OPN3; darker shades, devATR; lighter shades, sATR. Error bars, 95% CI. Note that the *OK371>OPN3* data are redrawn from Supplementary Figure 2, being the same flies with the same height and speed metrics, to enable direct graphical comparison.

**Supplementary Figure 8.**
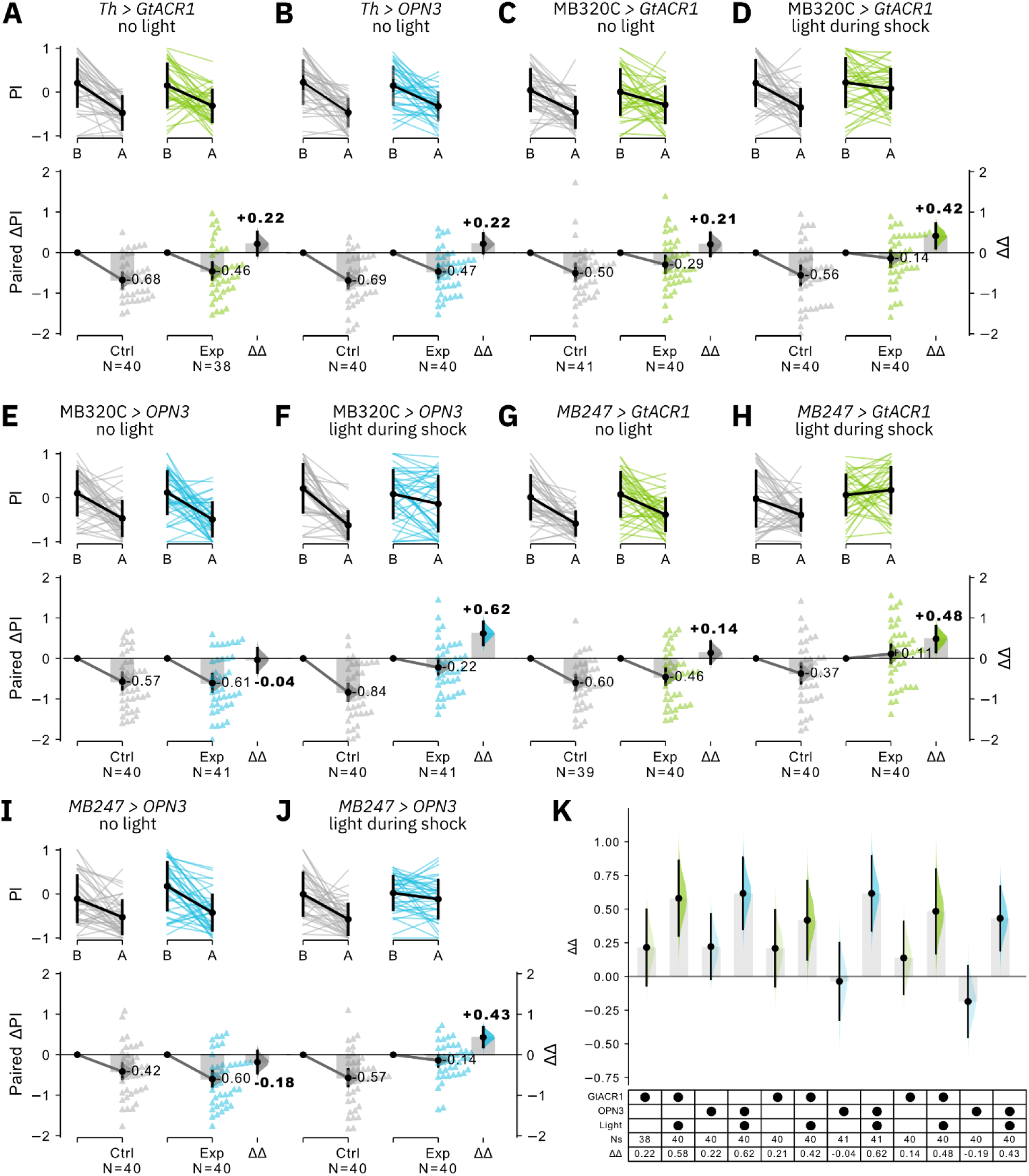
OPN3 successfully inhibits dopaminergic neurons and Kenyon cells during associative learning. **(A-B)** When no light is presented during the shock epoch, mild impairment of learning is observed in *Th>GtACR1* and *Th>OPN3* flies compared to genetic controls (i.e. pooled driver-only and responder-only controls). **(C-D)** Learning is slightly impaired in *MB320*>*GtACR1* flies in light-off conditions and significantly impaired in light-on conditions. **(E-F)** Learning is not impaired in *MB320>OPN3* in light-off conditions, but significantly impaired in light-on. **(G-H)** Learning is slightly impaired in *MB247*>*GtACR1* flies in light-off conditions and significantly impaired in light-on conditions. **(I-J)** Learning is not impaired in *MB247>OPN3* in light-off conditions, but significantly impaired in light-on. **(K)** Summary forest plot of ΔΔPIs from panels A-J and Figure 4C-D. Means and CIs for all panels are available in Supplementary Data.

**Supplementary Figure 9.**
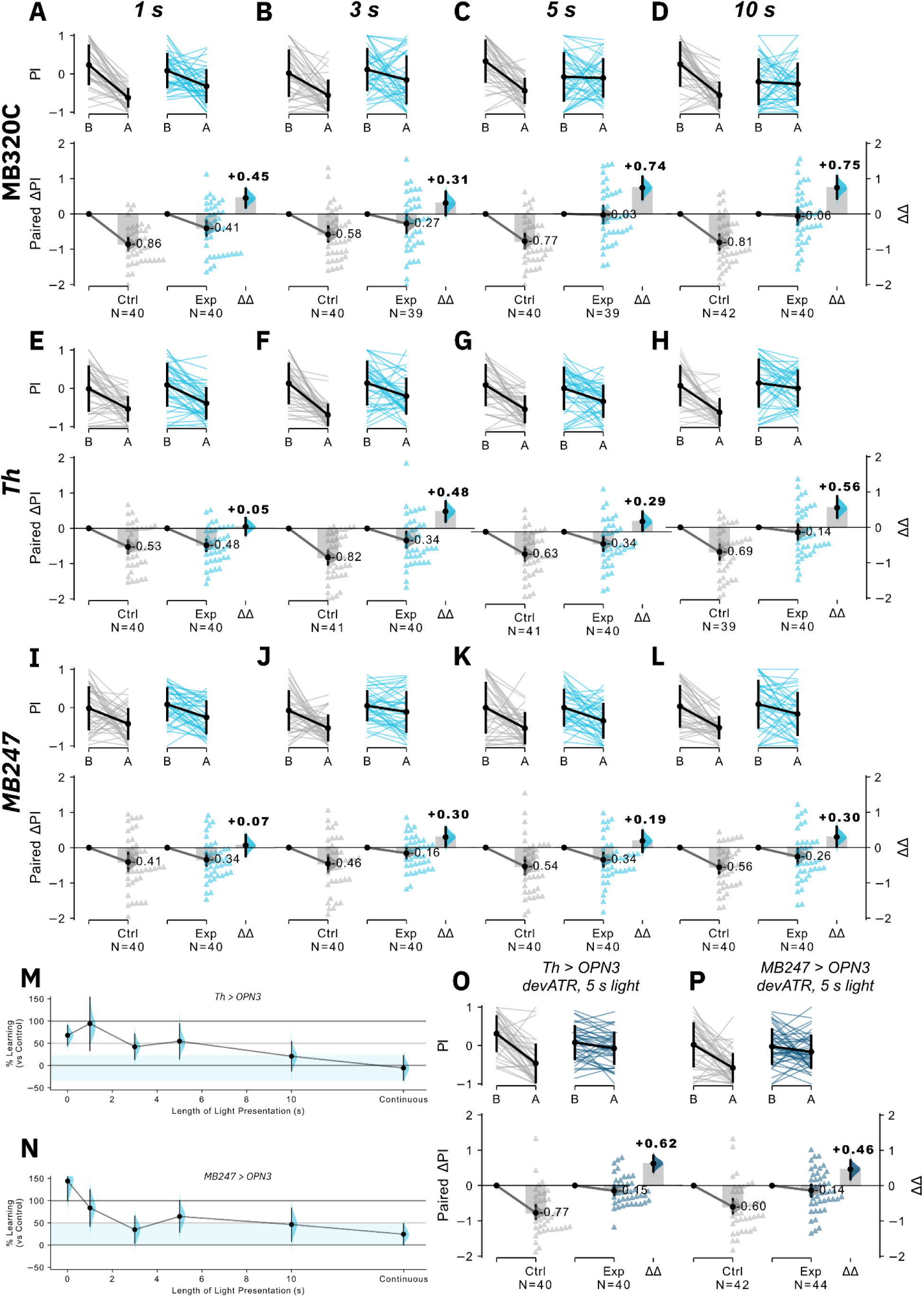
A short pulse of light induces long-term inhibition in dopaminergic neurons and Kenyon cells expressing OPN3. **(A-D)** *MB320C>OPN3* flies were presented with 1 s, 3 s, 5 s or 10 s prior to shock training. Mild impairment was observed with 1 s and 3 s stimulation, and complete ablation of learning at 5 s and 10 s. See also Figure 4G. **(E-H)** *Th>OPN3* flies were presented with 1 s, 3 s, 5 s or 10 s prior to shock training. Mild impairment was observed with 1 s, 3 s and 5 s stimulation, and near complete ablation of learning at 10 s. See also panel M. **(I-L)** *MB247>OPN3* flies were presented with 1 s, 3 s, 5 s or 10 s prior to shock training. Mild impairment was observed with 1 s, 3 s and 5 s stimulation, and near complete ablation of learning at 10 s. See also panel N. **(M)** Summary plot of % learning in *Th>OPN3* flies compared to genetic controls (pooled driver-only or responder-only controls) at 0 s, 1 s, 3 s, 5 s, 10 s, and continuous light. 0 s and continuous light data is the same as light-on and light-off data in Figure 4C-D. **(N)** Summary plot of % learning in *MB247>OPN3* flies compared to genetic controls (pooled driver-only or responder-only controls) at 0 s, 1 s, 3 s, 5 s, 10 s, and continuous light. 0 s and continuous light data is the same as light-on and light-off data in Supplementary Figure 8I-J. **(O)** *Th>OPN3* flies raised on ATR food show greater memory impairment with 5 s of light presentation compared to without (compare panel G). **(P)** *MB247>OPN3* flies raised on ATR food show greater memory impairment with 5 s of light presentation compared to without (compare panel K). Means and CIs for all panels are available in Supplementary Data.

**Supplementary Figure 10.**
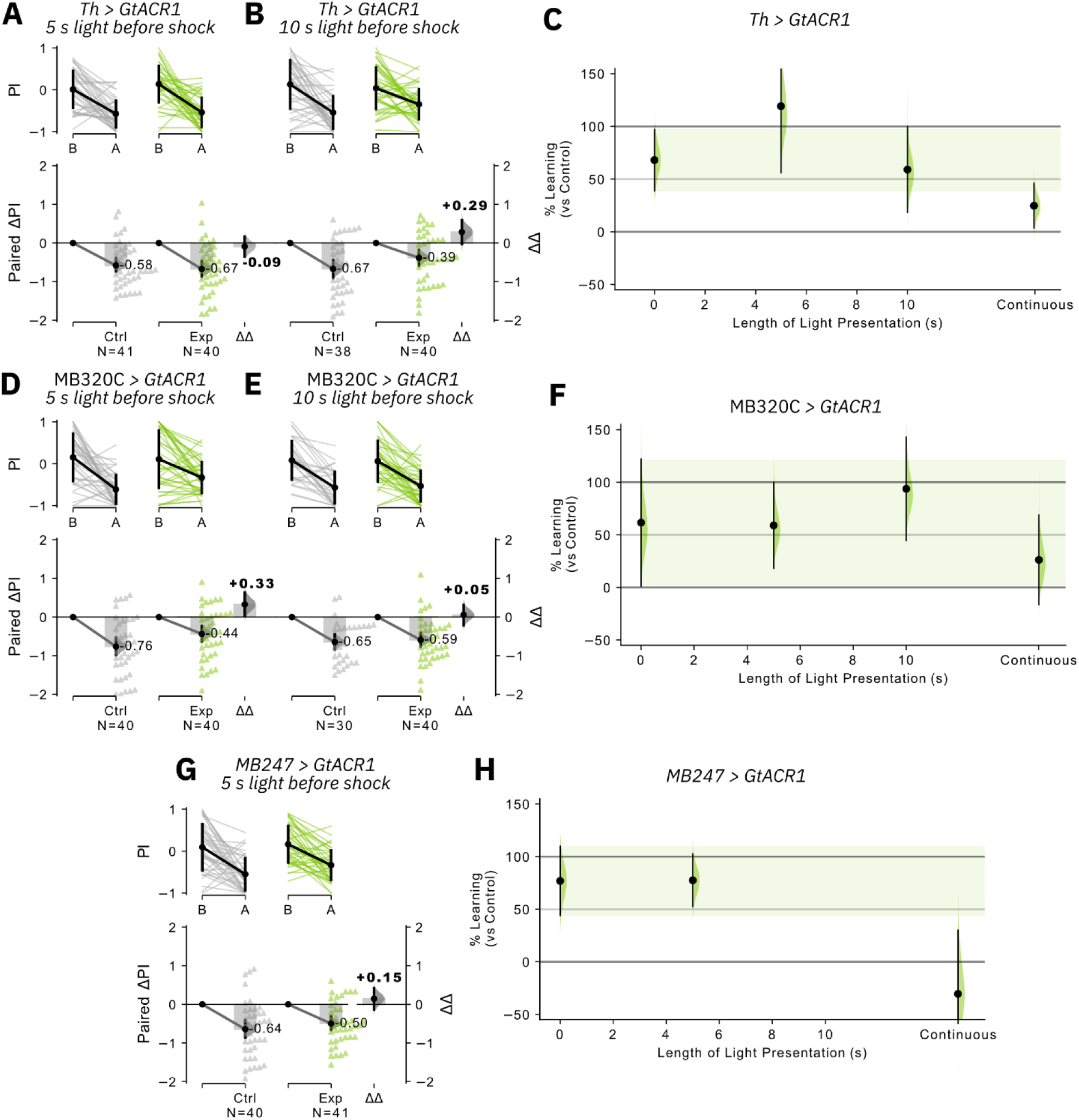
A short pulse of light does not induce long-term inhibition in dopaminergic neurons and Kenyon cells expressing GtACR1. (A-B) *Th>GtACR1* flies were presented with either 5 s or 10 s light before shock. Neither condition resulted in appreciable impairment of learning compared to no-light controls (compare Supplementary Figure 8A). **(C)** Summary of % learning of *Th>GtACR1* flies with 0 s, 5 s, 10 s or continuous light presentation. 5 s or 10 s light did not reduce learning below controls. Data for 0 s and continuous light are identical to Supplementary Figure 8A and Figure 4C respectively. Shaded region: 95% CI of light-off condition. **(D-E)** MB320C>*GtACR1* flies were presented with either 5 s or 10 s light before shock. Neither condition resulted in appreciable impairment of learning compared to no-light controls (compare Supplementary Figure 8C). **(F)** Summary of % learning of MB320C*>GtACR1* flies with 0 s, 5 s, 10 s or continuous light presentation. 5 s or 10 s light did not reduce learning below controls. Data for 0 s and continuous light are identical to Supplementary Figure 8C-D. Shaded region: 95% CI of light-off condition. **(G)** *MB247*>*GtACR1* flies were presented with 5 s light before shock, which did not reduce learning compared to controls (compare Supplementary Figure 8G). **(F)** Summary of % learning of *MB247>GtACR1* flies with 0 s, 5 s or continuous light presentation. 5 s light did not reduce learning below controls. Data for 0 s and continuous light are identical to Supplementary Figure 8G-H. Shaded region: 95% CI of light-off condition. Means and CIs for all panels are available in Supplementary Data.

**Supplementary Figure 11.**
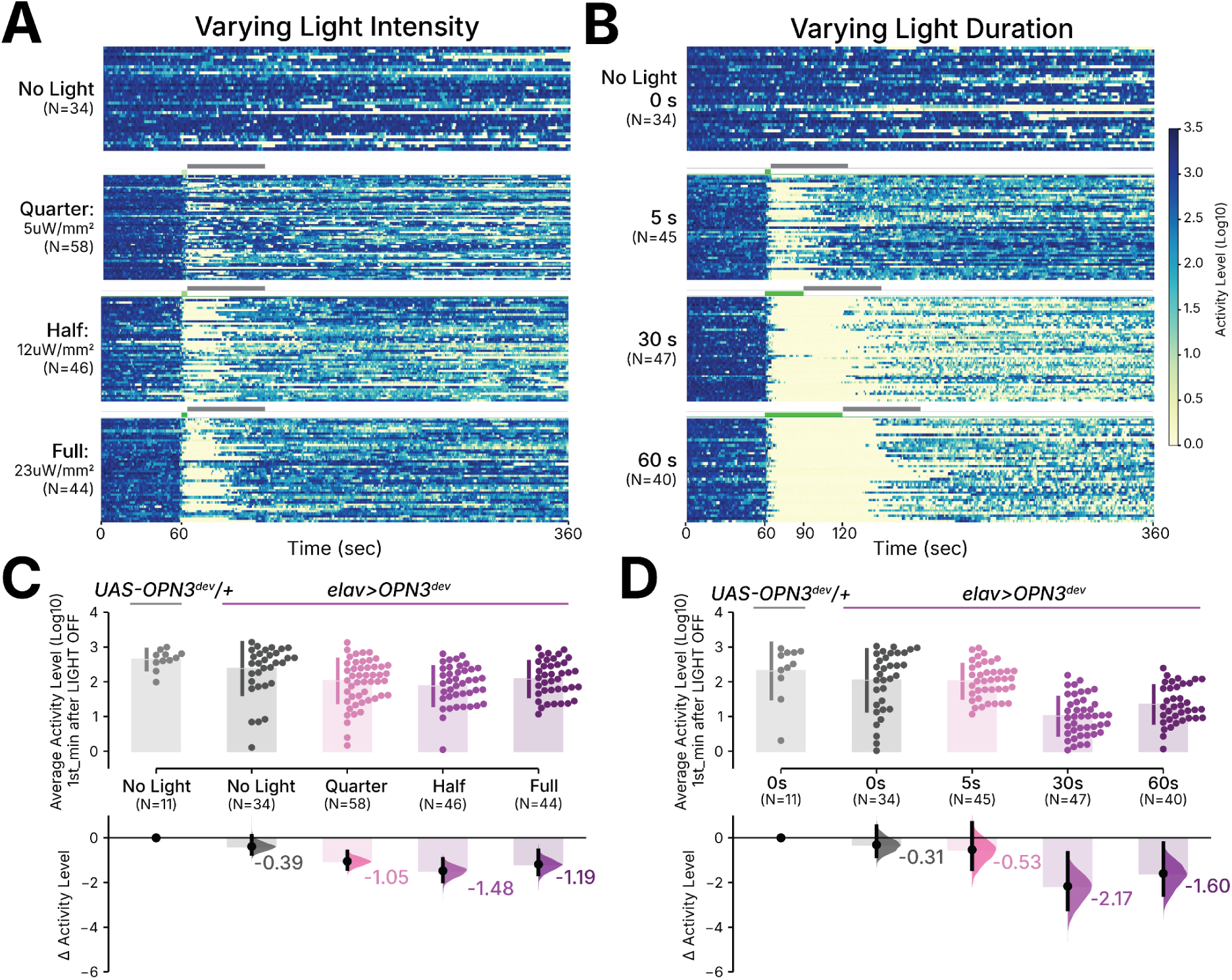
Activity Level plots for *elav>OPN3*^dev^ in the *Trumelan* assay. Individual fly activity heatmaps for *elav>OPN3*^dev^ flies tested under (**A**) varying light intensities (No Light, Quarter: 5 μW/mm^2^; Half: 12 μW/mm^2^; Full: 24 μW/mm^2^) and (**B**) light durations (0 s, 5 s, 30 s, 60 s). Each row represents a single fly. Average speed plots 1 min after light offset for UAS-*OPN3*^dev^*/+* controls and *elav>OPN3*^dev^ flies at (**C**) varying intensities and (**D**) durations (0 s, 5 s, 30 s, 60 s illumination period), with corresponding Hedge’s g effect sizes on the bottom with 95% confidence intervals. Superscripted ‘dev’ indicates flies were raised on ATR throughout development and post eclosion. N’s are sample sizes of each group. Note that the *elav>OPN3*^dev^ data are the same flies shown in Figure 5, with activity level presented here as an additional metric.

**Supplementary Figure 12.**
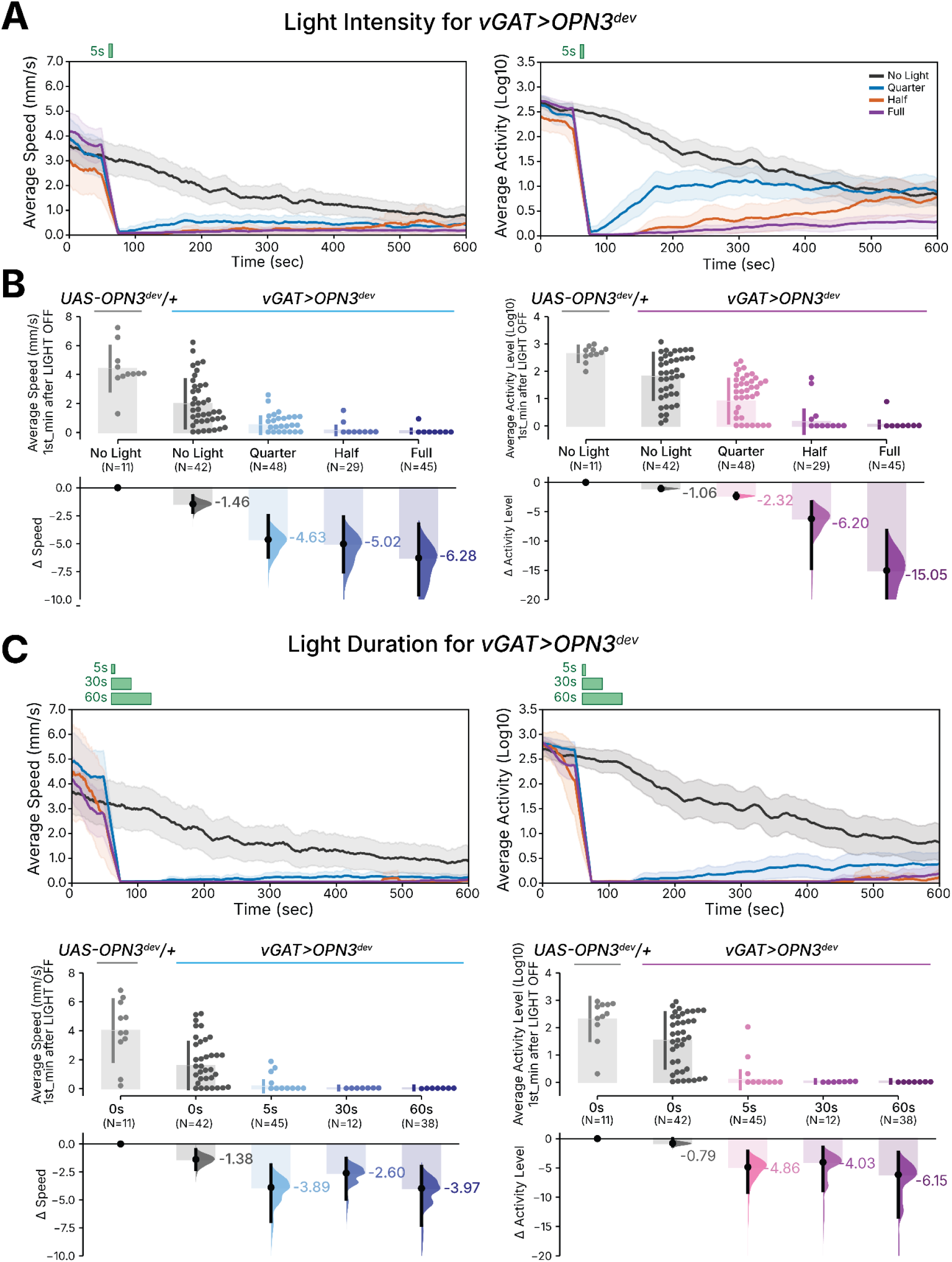
OPN3 produces robust prolonged silencing behavior when expressed in GABAergic neurons. (**A**) Average time-series traces of climbing speed (left) and activity level (right) for *vGAT > OPN3*^dev^ flies exposed to a brief 5 second green light pulse at varying intensities (No Light, Quarter: 5μW/mm^2^; Half: 12 μW/mm^2^; Full: 24 μW/mm^2^). (**B**) Average speed (left) and activity level (right) plots for light intensity 1 min after light offset for OPN3^dev^/+ controls and *vGAT>OPN3*^dev^. (**C**) Average time-series traces of climbing speed (left) and activity level (right) for *vGAT>OPN3*^dev^ flies exposed to a green light pulse at varying durations (0 s, 5 s, 30 s, 60 s illumination period). (**D**) Average speed (left) and activity level (right) plots for light duration 1 min after light offset for OPN3^dev^/+ controls and *vGAT> OPN3*^dev^. Green bar indicates illumination period. Superscripted ‘dev’ indicates flies were raised on ATR throughout development and post eclosion. Shaded regions represent 95% CI.

